# Distinct protocerebral neuropils associated with attractive and aversive female-produced odorants in the male moth brain

**DOI:** 10.1101/2020.12.11.421289

**Authors:** Jonas Hansen Kymre, XiaoLan Liu, Elena Ian, Christoffer Nerland Berge, XinCheng Zhao, GuiRong Wang, Bente G. Berg, Xi Chu

## Abstract

The pheromone system of heliothine moths is an optimal model for studying principles underlying higher-order olfactory processing. In *Helicoverpa armigera*, three male-specific glomeruli receive input about three female-produced signals, the primary pheromone component, serving as an attractant, and two minor constituents, serving a dual function, i.e. attraction versus inhibition of attraction. From the antennal-lobe glomeruli, the information is conveyed to higher olfactory centers, including the lateral protocerebrum, via three main paths – of which the medial tract is the most prominent. In this study, we traced physiologically identified medial-tract projection neurons from each of the three male-specific glomeruli with the aim of mapping their terminal branches in the lateral protocerebrum. Our data suggest that the neurons’ wide-spread projections are organized according to behavioral significance, including a spatial separation of signals representing attraction versus inhibition – however, with a unique capacity of switching behavioral consequence based on the amount of the minor components.

## Introduction

Olfactory circuits serve a central role in encoding and modulating sensory input from the natural surroundings. Understanding how these chemosensory circuits translate signals with different hedonic valences into behavior is an essential issue in neuroscience. With a relatively simple brain and a restricted number of associated odors evoking opposite innate behaviors, i.e. attraction and aversion, the insect pheromone pathway is an optimal system to address this question. In the noctuid moth, pheromone-evoked behaviors are linked to a hardwired circuit in the lateral protocerebrum, including the lateral horn. This brain region shares many neural principles with the mammalian amygdala (Miyamichi et al., 2011; Sosulski et al., 2011). In contrast to the random neuron connectivity in another higher-order olfactory center of the insect, the mushroom body calyx (mammalian piriform cortex analog, Su et al., 2009), the neuronal wiring in the lateral protocerebrum is characterized by a form of spatial clustering. Here, neurons responding to food odors versus pheromones as well as attractive versus aversive odors are spatially segregated (Grabe & Sachse, 2018). Thus, at the level of the lateral protocerebrum, it appears that the relevant odor cues are represented in different wide-spread sub-domains that display a form of spatial pattern according to behavioral significance.

In the moth antennal lobe (AL; mammalian olfactory bulb analog), the male-specific macroglomerular complex (MGC) receives input from olfactory sensory neurons (OSNs). The MGC glomeruli process input about a few female-produced signals, of which one primary constituent acts as an unambiguous attractant and others often enhance attraction at low doses but serves as behavioral antagonists at higher doses (Chang et al., 2017; Gothilf et al., 1978; Kehat & Dunkelblum, 1990; Wu et al., 2015; Zhang et al., 2012). From the MGC, the pheromone signals are conveyed to higher brain centers, including the lateral protocerebrum, by male-specific projection neurons (PNs) following three main tracts, i.e. the medial, mediolateral, and lateral antennal-lobe tract (mALT, mlALT and lALT, respectively, see Homberg et al., 1988; Lee et al., 2019). Although a considerable number of medial-tract MGC neurons have previously been reported in various moth species (Anton et al., 1997; Berg et al., 1998; Christensen & Hildebrand, 1987; Christensen et al., 1991; Christensen et al., 1995; Hansson et al., 1994; Hansson et al., 1991; Jarriault et al., 2009; Kanzaki et al., 1989; Kanzaki et al., 2003; Nirazawa et al., 2017; Seki et al., 2005; Vickers et al., 1998; Zhao & Berg, 2010; Zhao et al., 2014), the main focus has so far been odor coding within the MGC. Only a few studies have paid attention to the protocerebral projections of these medial-tract PNs (Kanzaki et al., 2003; Seki et al., 2005; Zhao et al., 2014). Thereby, our two main questions are where in the lateral protocerebrum these neurons project to and how the pheromone neural circuit at this level processes information with opposite valences.

The male moth used in this study, *Helicoverpa armigera* (Lepidoptera, *Noctuidae, Heliothinae*), utilizes *cis*-11-hexadecenal (Z11-16:Al) as the primary pheromone component and *cis-*9-hexadecenal (Z9-16:Al) as a secondary component (Kehat & Dunkelblum, 1990). Notably, the secondary constituent is identical with the primary pheromone component of the coresidential and closely related species, *Helicoverpa assulta* (Berg et al., 2014), and could thus act as an aversive signal at high concentrations. Another female-produced minor component, *cis*-9-tetradecenal (Z9-14:Al), undoubtedly plays a dual role, acting as a behavioral antagonist, i.e. inhibiting the attraction elicited by the primary pheromone at higher dosages, i.e. >5% (Gothilf et al., 1978; Kehat & Dunkelblum, 1990), and as an agonist at lower concentrations, i.e. 0.3-5% (Wu et al., 2015; Zhang et al., 2012). This functional duality of a single molecular component indicates the complexity of the neural circuits processing pheromone information. The system must be capable of encoding attractive and aversive signals appropriately in order to elicit coordinated responses maximizing reproductive fitness.

In the study presented here, we characterized the male-specific PNs passing along the prominent mALT and the slightly thinner mlALT in *H. armigera*, with focus on their projection patterns in the lateral protocerebrum. Based on the intracellular recording/staining technique, combined with calcium-imaging experiments, we have mapped the projection patterns of physiologically identified MGC-neurons. The current study provides solid evidence for a spatial arrangement within the relevant protocerebral neuropils of the male moth demonstrating distinct regions receiving input about separate or intermixed female-released signals associated with attraction versus inhibition of attraction.

## Results

### Mapping odor-evoked responses of the MGC output neurons by means of calcium imaging

The projection pattern of the male-specific *sensory* neurons onto the MGC units was previously mapped via bath application calcium-imaging studies (Kuebler et al., 1998; Wu et al., 2013; Wu et al., 2015). To measure the output signals from the same three MGC glomeruli, we performed calcium-imaging measurements of odor-evoked responses in a group of PNs exclusively. By applying a calcium-sensitive dye (Fura 2) into the calyces (Fig. 1A), we label primarily the population of medial-tract uniglomerular neurons (Ian et al., 2016; Kymre et al., 2020). In the species used here, *H. armigera*, the MGC comprises three units (Skiri et al., 2005; Zhao et al., 2016) receiving input from three OSN categories: (1) the cumulus from OSNs tuned to the primary component, Z11-16:Al, (2) the dorsomedial posterior (dmp) unit from OSNs tuned mainly to the secondary component, Z9-16:Al, and (3) the dorsomedial anterior (dma) unit from OSNs tuned to the behavioral antagonist/enhancer, Z9-14:AL (Wu et al., 2015). For simplicity, Z9-14:Al is mentioned as a behavioral antagonist in the subsequent text. The imaging data of mALT PNs connected to each of the three MGC units was obtained from dorsally oriented brains (Fig. 1B). One example of repeated traces illustrated that the increase in intracellular Ca^2+^ in the MGC during antennal stimulation with the two pheromone components was consistent (Fig. 1C).

**Figure 1.**
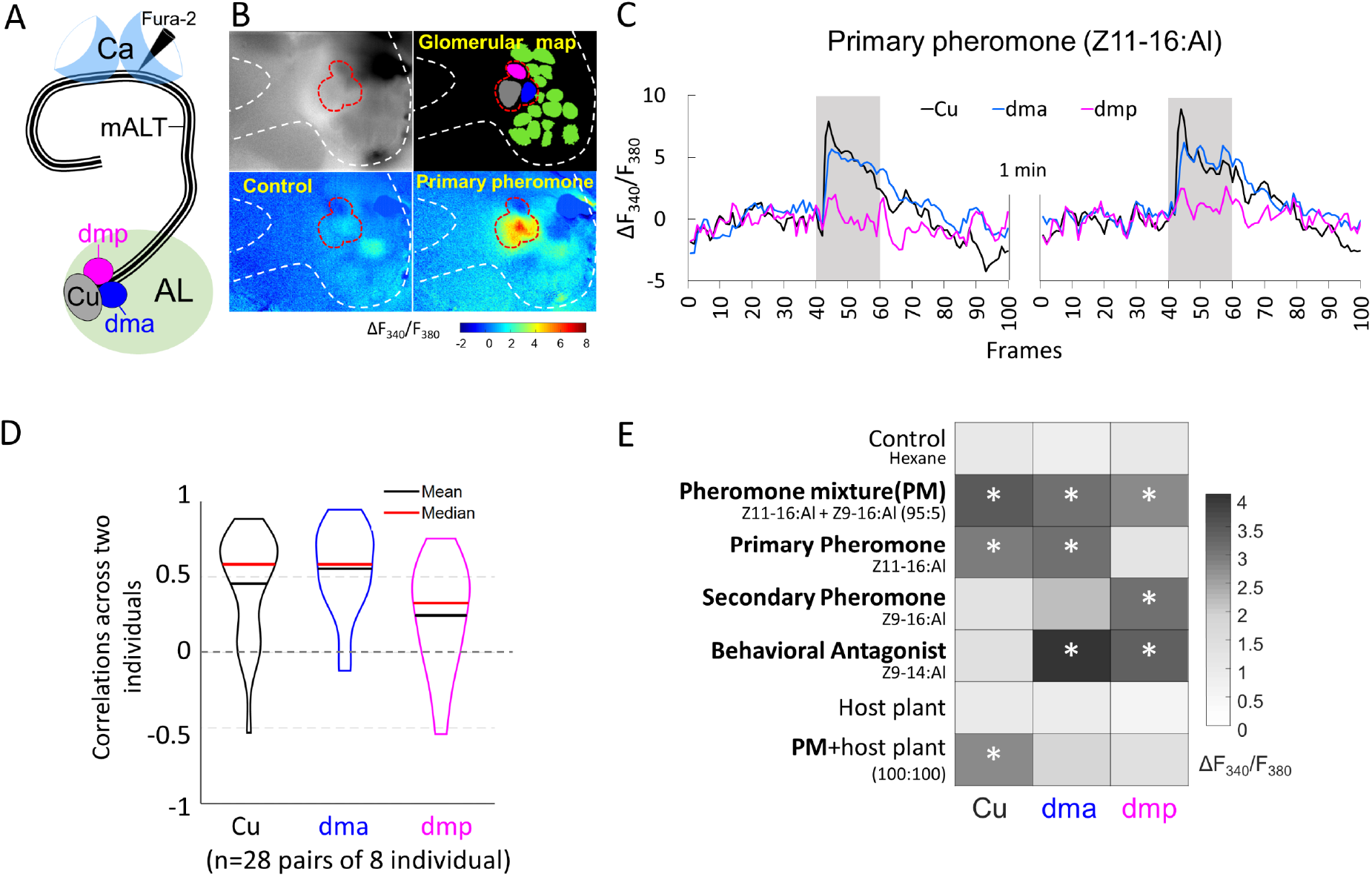
MGC neurons confined to the media tract (mALT) and their odor responses during calcium imaging. (**A**) Illustration of the retrograde staining from calyx (Ca) labeling the MGC mALT neurons. (**B**) Pictorial material representing calcium imaging data: top-left, raw image of an AL stained with Fura from the Ca; top-right, a processed image showing a map of recognized glomeruli; down-left and down-right, Heat maps of responses to the control and primary pheromone. White dash border circumvents the area of AL, red border shows the MGC region. (**C**) Example of calcium imaging traces showing response to the primary pheromone from neurons innervating distinct MGC units. The standardized traces quantify the neuronal activity of two repeated stimulations with 100 ms sampling frequency. The interval between stimulations is 1 min. Gray bar, the duration of the stimulus (2 s). (**D**) Violin plot of consistent tests across 8 individuals. (**E**) Response amplitudes of a population of PNs innervating the same MGC units to all presented stimuli, where * indicates a significant response compared with control.

We also checked whether the population of mALT PNs from each MGC unit showed consistent responses across individual insects. The response consistency of a group of PNs can be quantified as the Pearson’s correlation coefficient of the response vectors in two individual insects, where each vector contains the trial-averaged responses of an individual to a given set of odors (Mittal et al., 2020; Schaffer et al., 2018). The average correlation between the cumulus medial-tract PNs responses (mean calcium signal during stimulation windows subtracted by that during the pre-stimulation period) across individuals was 0.45. The corresponding average correlations of PNs from the dma and dmp units were 0.55 and 0.24, respectively (Fig. 1D). These paired-individual correlations were greater than the nonlinear relationship with a chance level of 0 (*t*-test, *p*s<0.002), confirming the general response consistency across different insects. We profiled the pheromone responses in mALT PNs from each MGC unit in 8 insects, by comparing the Fura signals evoked by each stimulus during the 2s stimulation window with the control (hexane) (Fig. 1E). The average calcium traces observed in these insects are presented in Figure 1 – figure supplement 1. As expected, the cumulus PN population showed a pronounced activation during stimulation with the primary pheromone and the pheromone blend. The dma output neuron population, in turn, responded not only to the behavioral antagonist but also to the primary pheromone and the pheromone blend. The dmp projection neurons responded to the secondary pheromone and the behavioral antagonist, as well as to the pheromone mixture. All these responses showed a phasic component that decayed over the course of the stimulation period. Taken together, each stimulus evoked a unique activation pattern in the three MGC units (Fig. 1E). We also analyzed the mean calcium trace of 8 individuals across 7 stimuli (see *Odor stimulation* section in Materials and methods). The across-stimuli correlation plot illustrates that, unlike the relatively defined responses of the PNs innervating the cumulus and the dmp unit, the population of dma PNs evoked a broad activation pattern including responses to all female-produced chemicals tested (Figure 1 – figure supplement 2). In contrast to the narrowly tuned male-specific OSNs, we found that the odor response profiles of the medial-tract output neurons were considerably more intricate.

### Morphological characteristics of individual MGC projection neurons

We next aimed to elucidate the functioning and morphology of individual neurons involved in processing pheromone information. Intracellular recording and staining were executed from the thick dendrites of MGC output neurons, including PNs confined to both the mALT and mlALT (Fig. 2 and Fig. 2 – figure supplement 1). We labeled 35 PNs across 32 preparations (see supplementary table S1), along with four preparations with multiple co-labeled heteromorphic MGC neurons.

**Figure 2.**
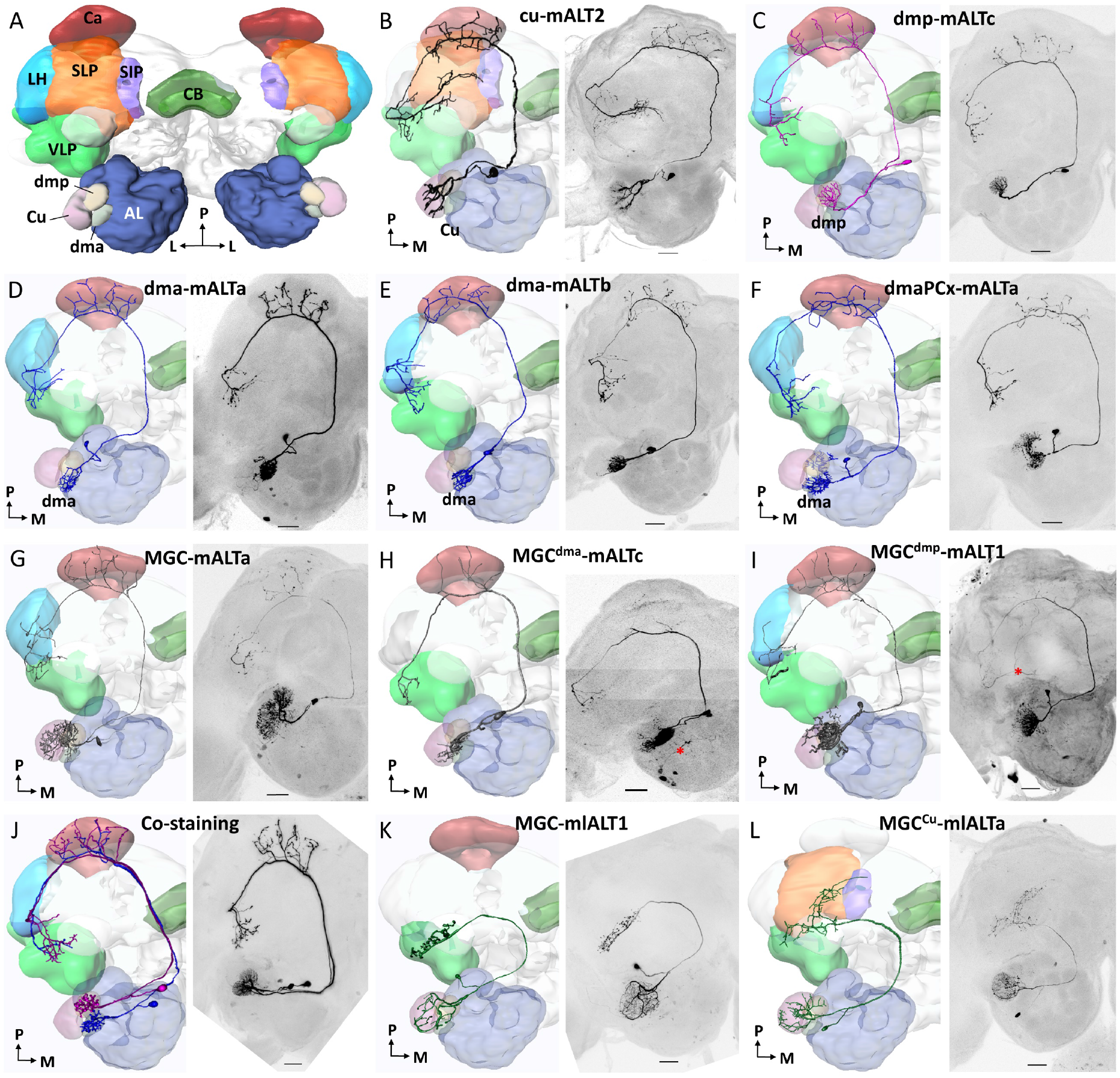
Morphological features of PNs in the medial and mediolateral tracts - reconstruction and confocal images. (**A**) Diagram of the brain neuropils targeted by the MGC output neurons in a dorsal view. Color codes are in correspondence with all other figure panels. AL, antennal lobe; Ca, calyces; CB, central body; LH, lateral horn; SIP, superior intermediate protocerebrum; SLP, superior lateral protocerebrum; VLP, ventrolateral protocerebrum. (**B-E**) Examples of uniglomerular PNs in the mALT, originating from each of the MGC units. One of two morphological types of dma mALT PNs in (**D**), with terminals in the Ca, anteroventral LH, VLP and approaching SLP. The other dma PN type (**E**) is more restricted to LH and VLP. (**F)** A multiglomerular PN in mALT with dendritic innervation in dma and three posterior complex (PCx) glomeruli. (**G-I**) Multiglomerular MGC PNs, with dendritic branches innervating the MGC units homogeneously (**G**), predominantly in the dma (**H**), or mainly in the dmp (**I**). (**J**) Two co-labeled mALT PNs, one innervating the dma (blue) and another arborizing in the dmp (magenta). Both PNs sent their axon terminals to overlapping regions of the LH and VLP. (**K**) A mlALT PN with homogeneously distributed dendrites across all MGC units. (**L**) Another type of mlALT PN, with dendrites in all MGC units, but dense arborizations only in the cumulus. Unique neuron IDs are presented. All 3D reconstructions were manually registered into the representative brain. D, dorsal; L, lateral; M, medial; P, posterior. Red asterisks indicate weakly co-labeled neurons. Scale bars: 50 μm.

#### MGC PNs in the medial tract

In total, 29 individual MGC PNs confined to the mALT were stained. For simplification, any identical PNs that were co-labeled are referred to as a single PN, as physiological recordings contained only one analyzed waveform. All mALT PNs had their somata in the medial cell body cluster. Amongst these neurons, 24 had uniglomerular dendrites, of which 16 arborized in the cumulus (Fig. 2B), four in the dmp (Fig. 2C), and four in the dma (Fig. 2D-E). In addition, five mALT PNs had multiglomerular dendrites. One of these multiglomerular PNs arborized in both the dma and posterior-complex glomeruli (Fig. 2F), while the remaining four PNs innervated several MGC units, with different densities across the glomeruli. Two of these multiglomerular PNs had evenly distributed dendrites across the MGC (Fig. 2G), while the other two had dense innervation of either the dma or dmp, and sparse innervation in the remaining MGC glomeruli (Fig. 2H-I, respectively).

Generally, the protocerebral projection patterns of mALT PNs innervating the cumulus were homogeneous. These PNs projected sparsely to a restricted area in the inner layer of the calycal cups before targeting two main regions in the lateral protocerebrum, i.e. the ventrolateral protocerebrum (VLP) and superior lateral protocerebrum (SLP). Generally, the SLP was innervated through two separate branches targeting anterior and intermediate regions, respectively, with the most ventral parts positioned close to the superior clamp. In addition, at least seven of the 16 cumulus PNs innervated the posterodorsal part of the superior intermediate protocerebrum (SIP), localized adjacent to the vertical lobe. Besides the numerous cumulus-neurons, eight uniglomerular PNs originated in the two smaller MGC-units, four in dma and four in dmp. These PNs targeted primarily the VLP and the anteroventral lateral horn (LH), where they overlapped, as demonstrated from two preparations where uniglomerular dma- and dmp-PNs were co-labeled (Fig. 2J). Moreover, two dma-PNs had additional axon terminals in the region between the LH and SLP. The target areas of the five multiglomerular mALT PNs resembled those of the uniglomerular dma- and dmp-PNs, as their output terminals were confined to the VLP and anteroventral LH. All the mALT PN innervations in the VLP were spatially restricted, primarily into the most dorsoanterior regions of this neuropil.

#### MGC PNs in the mediolateral tract

In addition to the medial-tract PNs, we labeled three multiglomerular MGC PNs projecting through the mediolateral ALT, which is considerably thinner than the medial tract and projects directly to the lateral protocerebrum without innervating the calyces. Two of these PNs arborized in all three MGC units with evenly distributed dendrites (Fig. 2K). In contrast to most non-MGC mlALT PNs, innervating dorsal neuropils such as the SLP (Kymre et al., 2020), both PNs projected solely to ventral neuropils, mainly the VLP. Similar neurons were previously reported in another heliothine species (Lee et al., 2019). The final mlALT MGC-PN had dense dendritic arborizations in the cumulus and sparse innervations in the dma and dmp along with a few posterior-complex glomeruli (Fig. 2L). Its axonal projections targeted the VLP, SLP, and posterodorsal SIP, like the cumulus-PNs of the medial tract.

#### Output regions of PNs across distinct antennal-lobe tracts

Previous and present data including confocal images and 3D reconstructions of MGC PNs confined to the medial, mediolateral, and lateral ALTs, indicated that their output regions could potentially overlap in specific protocerebral regions (Chu et al., 2020a). To explore whether such PNs intersect, we next performed multi-staining experiments. One preparation contained three individual MGC neurons confined to each of the three main tracts (Fig. 3A). Here, one mALT PN and one lALT PN were strongly labeled, whereas the mlALT PN was moderately stained. The lateral-tract neuron projected to the ipsilateral column of the SIP, as previously described by Chu et al. (2020a). This region was not innervated by any other MGC-PN types. PNs confined to the medial and mediolateral tract, however, overlapped in the VLP (dashed circle in Fig. 3A). Two co-stained cumulus-PNs in another preparation, included one lateral-tract PN targeting the anteriorly located column of the SIP and one medial-tract PN targeting the VLP, SLP, and posterior SIP (Fig. 3B). These PNs had no overlapping terminals.

**Figure 3.**
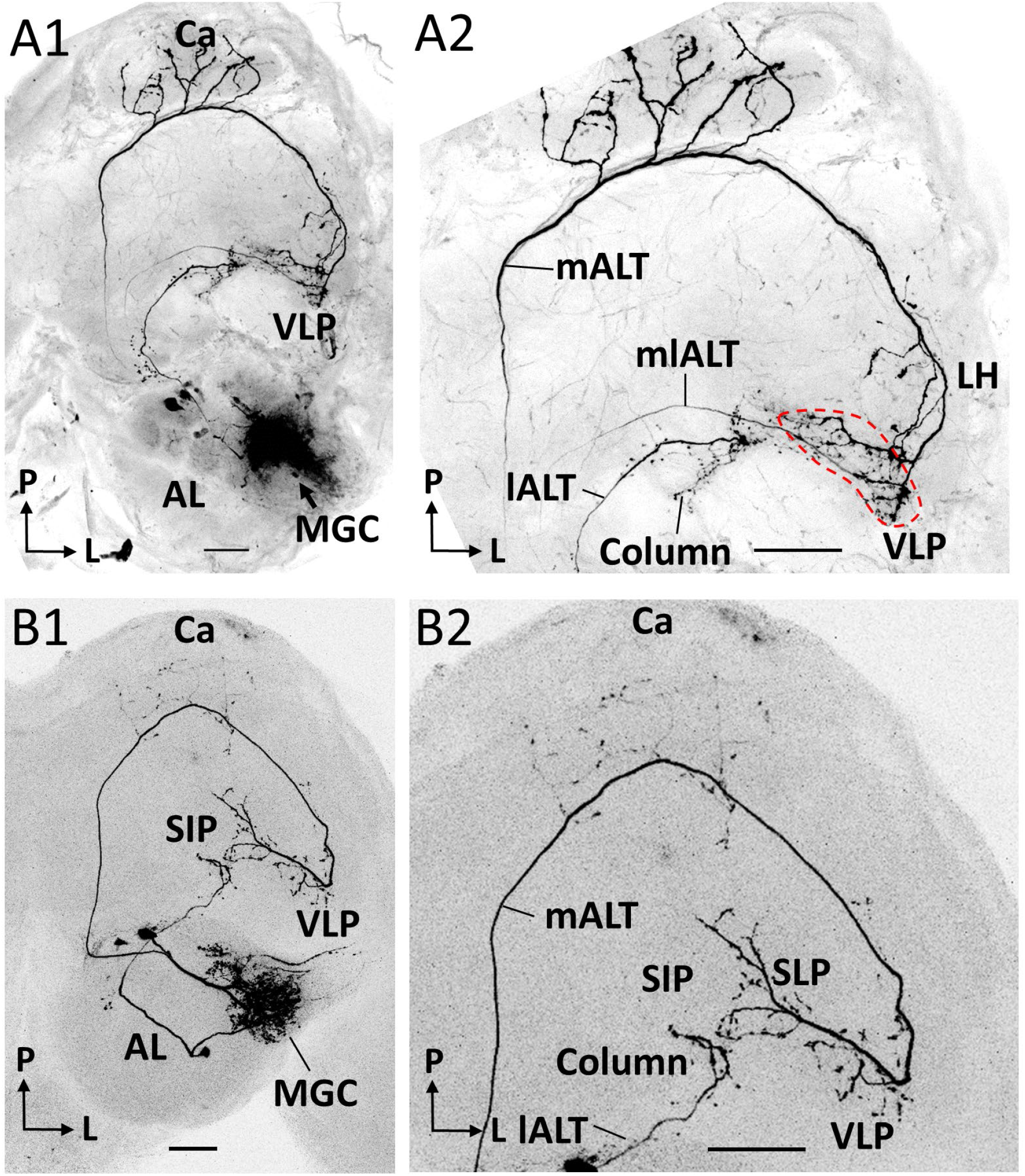
Co-labeled PNs in distinct ALTs. (**A**) Application of dye in the MGC visualized three antennal-lobe projection neurons confined to the mALT, mlALT and lALT, respectively (**A1**). The axonal terminals of the mlALT PN overlapped with the mALT PN in the VLP (red dashed lines in **A2**). (**B**) Two co-labeled cumulus PNs, confined to the mALT and lALT, respectively. The mALT PN has no overlap with the lALT PN. L, lateral; P, posterior. Scale bars: 50 μm.

### Physiological characteristics of the MGC PNs

Next, we characterized the electrophysiological features of the labelled PNs. The summarized mean spiking activity of all individual PNs, indicating which responses were significantly different from pre-stimulation firing rates, are reported in Fig. 4 (electrophysiological traces are in Fig. 4 - figure supplement 1–7). In addition, we report a heat map of every neuron’s mean Z-scored instantaneous firing rate during application of each female-produced stimulus (Fig. 5), as well as repeated trials of the same stimulus in individual PNs (Fig. 5 – figure supplement 1–3). Each neuron was named with a unique ID. For PNs having dendritic innervation only in one of the MGC units, the neuron ID is expressed as “innervated glomerulus-ALT number/letter”, and for PNs innervating multiple MGC units as “MGC^main innervated glomerulus^-ALT number/letter”. Here the field of “number/letter” represents two different concentration protocols (see *Odor stimulation* section in the Materials and methods). Finally, we averaged the responses of all individually recorded PNs, and describe how the sampled MGC PNs represent pheromone signals across distinct MGC units and protocerebral neuropils.

**Figure 4.**
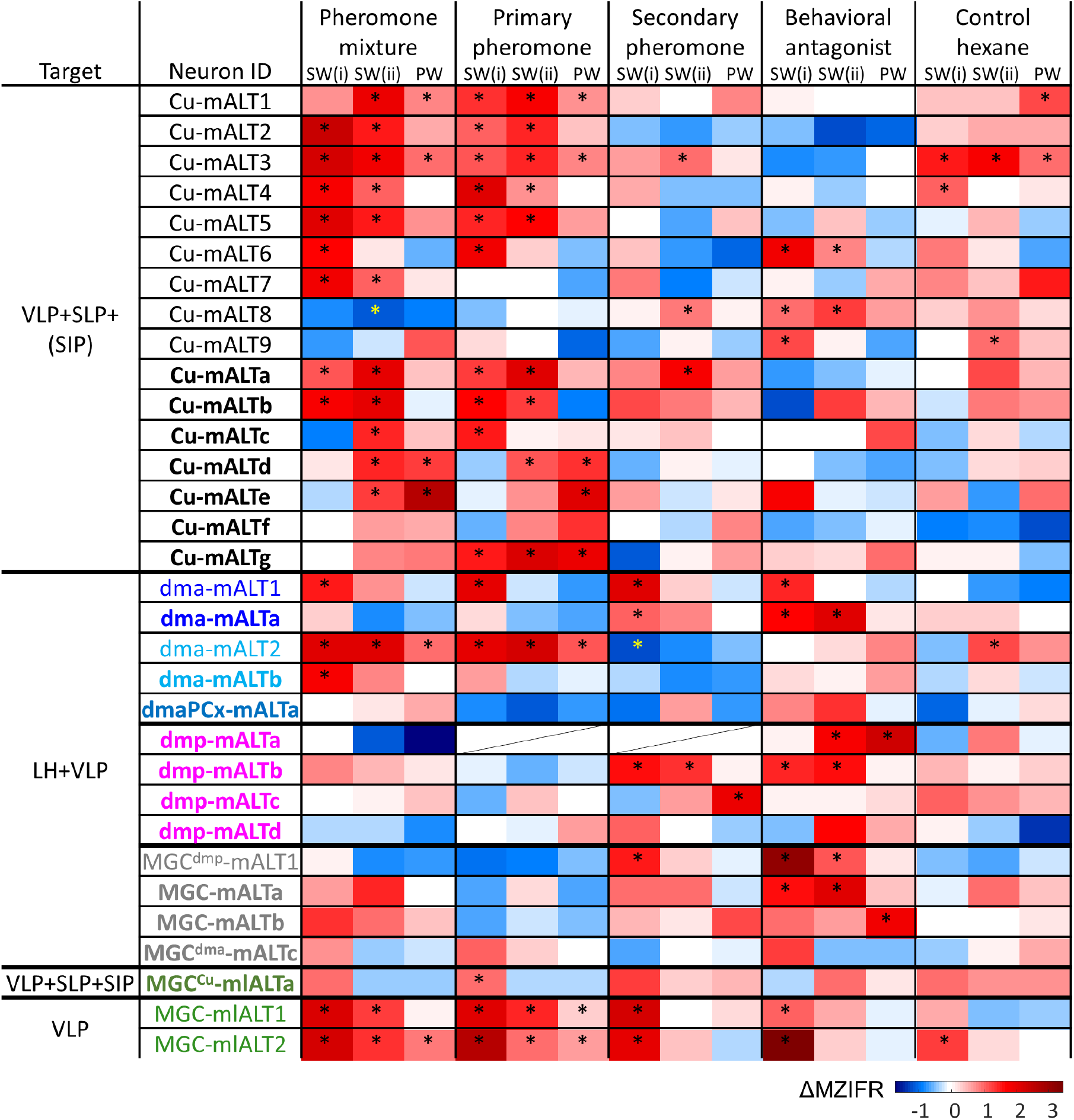
Physiological properties of individual MGC PNs. Mean response amplitudes (ΔMZIFR) were registered in three adjacent episodes: Sub-window SW(i) including the first 100 ms of the stimulation window, sub-window SW(ii) including the remaining 300 ms of the stimulus duration; and a 200 ms post-stimulation window (PW). *indicates significant response, determined according to the threshold of baseline activity of individual neurons (with 95% confidence level).

**Figure 5.**
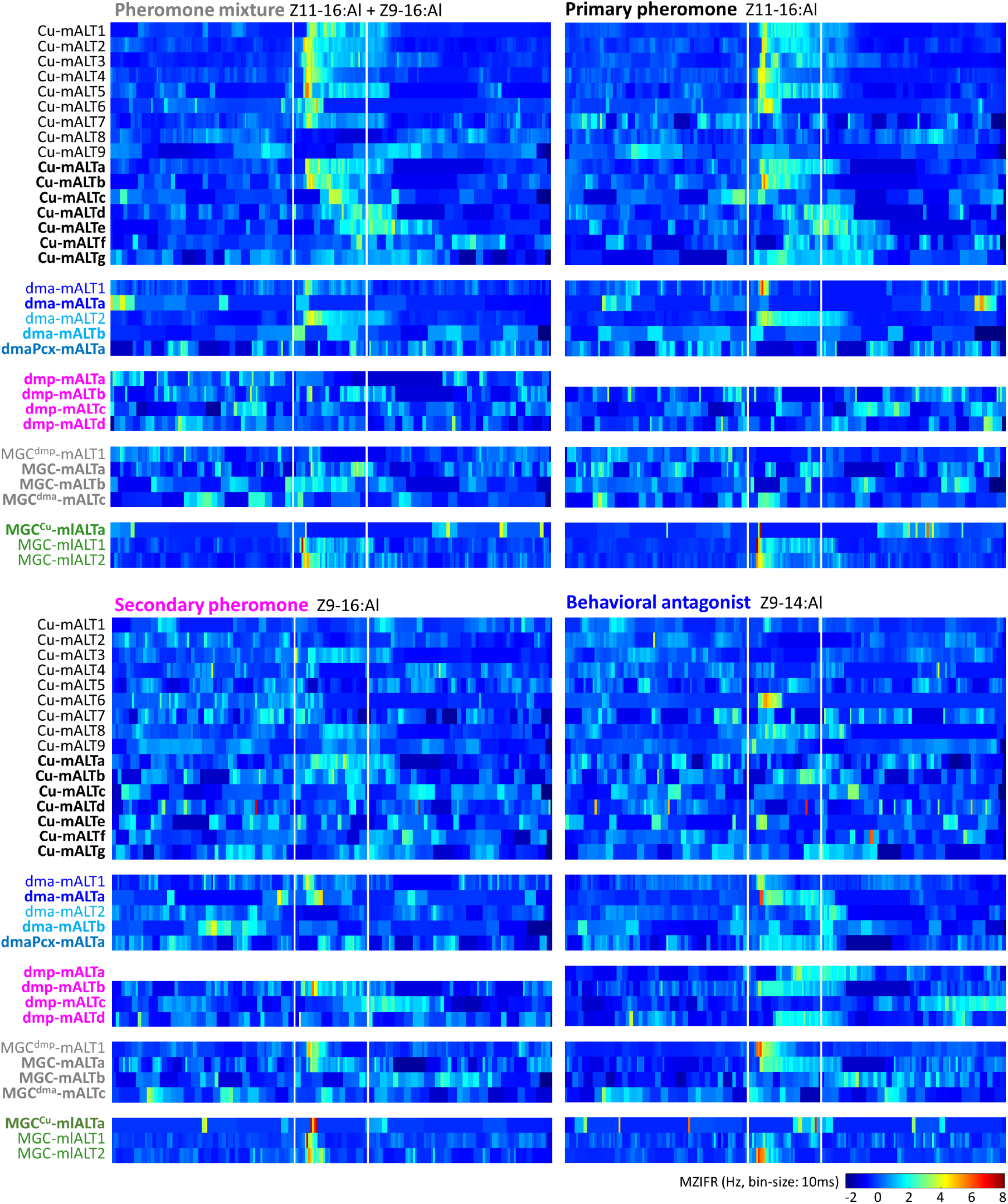
Temporal resolution index of individual MGC PNs. This plot displays the across-trials mean instantaneous firing rates (MZIFR) for each of the reported PNs, in response to the four presented female-produced stimuli.

#### Medial-tract PN response-profiles were only partly congruent with OSN inputs

The physiological profiles within each neuronal group innervating the same MGC unit varied to some extent, both with respect to odor discrimination and temporal response characteristics. The uniglomerular cumulus mALT PNs commonly responded with increased spiking frequency to both low and high concentrations of the primary pheromone and the pheromone mixture (Fig. 4 and Fig. 4 - figure supplements 1–2). In most cases, the responses were characterized by a sharp phasic onset that gradually faded towards baseline activity (Fig. 5). The phasic onset was most prominent for PNs in the low concentration protocol (Fig. 4 - figure supplement 1D); these neurons also appeared to discriminate between odorants more precisely than PNs in the high concentration protocol (Fig. 4 - figure supplement 1E). This coincides with previous studies reporting that OSNs display less specific response profiles with increasing odor concentrations (Malnic et al., 1999; Sato et al., 1994).

Notably, 25 % of the cumulus-PNs did not respond to the primary pheromone, and 31 % were excited by the behavioral antagonist and/or the secondary component (Fig. 4 and Fig. 4 – figure supplement 2). As these responses seem unlikely to arise due to OSN input, we hypothesized they might be mediated by local interneurons or multiglomerular MGC-PNs, which in case should imply delayed responses. Therefore, we quantified the onset and peak response latency of each recorded trial for the neurons responding to the relevant stimuli (Fig. 4 – figure supplement 2G). There were no clear distinctions across the different stimuli with respect to response *onset* latencies, but the *peak* latencies appeared to be delayed in response to the secondary pheromone.

The four uniglomerular dma PNs included two morphological sub-groups. The first one, including two PNs having fine branches approaching the SLP in addition to the terminals in the LH and VLP, responded with significant excitation to the behavioral antagonist and the secondary pheromone (Fig. 4 – figure supplement 3A-C). The second sub-group, consisting of the two remaining dma-PNs with restricted projections in the VLP and LH, were not excited by the behavioral antagonist nor the secondary compound (Fig. 4 and Fig. 4 – figure supplement 3D-F). Three of the dma PNs, including the two latter neurons, were excited by the pheromone mixture and/or primary pheromone. This is in agreement with our calcium-imaging data, demonstrating excitation in the population of dma-PNs during application of the stimuli including the primary pheromone. In addition to the uniglomerular dma-PNs, it is relevant to mention one multiglomerular PN here, arborizing in the dma and three posterior-complex glomeruli; this PN appeared to be excited by the behavioral antagonist and inhibited by the primary pheromone (Fig. 4 – figure supplement 4), but nonsignificantly (Fig. 4).

The four uniglomerular mALT PNs innervating the dmp were stimulated with the high-concentration protocol only. Even though the dmp unit reportedly receives input about the secondary pheromone and the behavioral antagonist (Wu et al., 2015), only one of the dmp-PNs identified here was significantly excited by both of these components (Fig. 4 – figure supplement 5). Of the three remaining dmp-PNs, one was excited by the behavioral antagonist exclusively, whereas two displayed no responses during any stimulus application.

The AL innervations of the four multiglomerular medial-tract PNs innervating all MGC units varied considerably. To investigate a putative association between MGC innervation and response characteristics, we quantified the dendritic density of each PN by measuring the fluorescence intensities within the separate MGC units (Fig. 4 – figure supplement 6), as previously performed (see Chu et al., 2020b; KC et al., 2020). One PN, arborizing primarily in the dmp, displayed phasic responses to the secondary pheromone and the behavioral antagonist. The second multiglomerular PN, innervating primarily the dma, had no responses to any of the female-produced compounds. The remaining two multiglomerular PNs, with approximately evenly distributed dendrites, had divergent response profiles. One exhibited a significant tonic excitation only to the behavioral antagonist, while the other showed no responses to any stimuli.

#### The responses of mlALT PNs corresponded with OSN inputs

The three mediolateral-tract MGC-PNs consisted of two morphological sub-types displaying different response profiles. The singular PN constituting the first sub-type targeted the VLP, SLP, and SIP, while its dendrites filled the cumulus densely and the dma, dmp, and some posterior complex glomeruli sparsely. This neuron responded with a weak and early phasic excitation to most stimuli. Yet, the primary pheromone induced the only significant response (Fig. 4 & 5; Fig. 4 – figure supplement 7). The other sub-type, including two mlALT PNs, covered the MGC units quite uniformly. These PNs were more broadly tuned, responding with excitation to all stimuli, however their responses to the pheromone mixture and the primary pheromone lasted substantially longer than the phasic excitations elicited by the secondary component and the behavioral antagonist (Fig. 4 – figure supplement 7; Fig. 5).

#### Representation of output signals from MGC PNs

Based on the precise electrophysiological and morphological data obtained from intracellular recordings, we took the mean response of MGC medial- and mediolateral-tract PNs, to provide a summary of odor-stimuli representation across the separate MGC units and the protocerebral output regions, respectively (Fig. 6). We first categorized the neurons into four groups based on their dendritic arborization, i.e. in the cumulus, dma, dmp, or in all MGC units. The mean responses showed that mALT PNs with dendrites in the cumulus deal with the primary pheromone, and those in the dmp unit with the secondary pheromone and the behavioral antagonist (Fig. 6A). This is consistent with the calcium-imaging results on medial-tract PNs (Fig. 1), and with former reports on response properties of the corresponding OSNs (Wu et al., 2015). However, unlike previous reports on OSN input, both our individual PN recordings and calcium-imaging tests indicated a role for the dma PNs in processing information not only about the behavioral antagonist, but also, surprisingly, about the primary pheromone and pheromone mixture. However, averaging the activity across the uniglomerular dma neurons, demonstrated that the most potent stimuli for these PNs were the behavioral antagonist and the pheromone mixture, while the primary pheromone elicited only minor increases in mean firing rates. The responses to the mixture and the primary pheromone could be influenced by other multiglomerular antennal-lobe neurons. For the fourth group, which consists of the multiglomerular mALT and mlALT PNs arborizing in the entire MGC, the averaged data demonstrated activation by all female-produced components.

**Figure 6.**
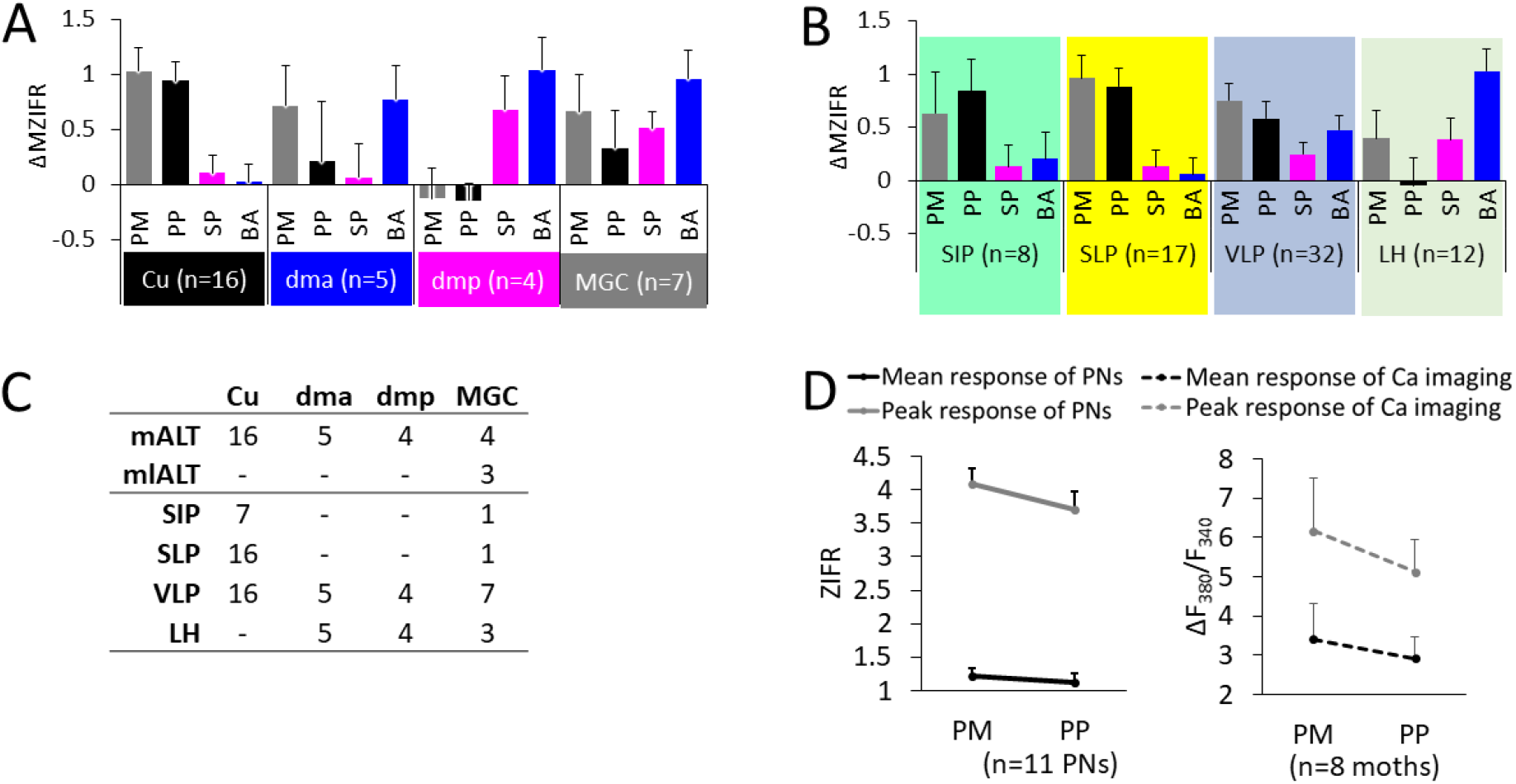
Summary of odor representation across MGC units and protocerebral neuropils. (**A**) Mean responses (ΔMZIFR; delta mean Z-scored instantaneous firing rate, for details see *Spike data analysis* in *Materials and methods*) of MGC medial/mediolateral tract PNs in response to distinct female-produced odorants, sorted according to the dendritic arborization. Data is presented in mean+se. Note that the PNs listed as innervating the cumulus (Cu), dma, and dmp were uniglomerular, while the PNs in the MGC category were multiglomerular. (**B**) Mean responses sorted by the protocerebral projections of the relevant PNs. The SIP and SLP predominantly receive input regarding attraction, while the aversive signals are strongly overrepresented in the anteroventral LH. The VLP receives signals regarding all tested pheromone-related stimuli. (**C**) Overview of all PNs included in **A** & **B**. As the PNs commonly innervated more than one protocerebral neuropil, individual PNs are represented in several neuropils in **B**. (**D**) Mean responses of pheromone mixture and primary pheromone in individual PNs (left) and in calcium imaging tests (right). Data is presented in mean+se. PM, pheromone mixture; PP, primary pheromone, SP, secondary pheromone; BA, behavioral antagonist.

Next, we investigated how the activity of MGC medial- and mediolateral-tract PNs is represented in the higher brain regions (Fig. 6B). For the protocerebral output areas, several neuropils, such the VLP, LH, SLP, and the posterior SIP, contained terminals of medial/mediolateral-tract MGC neurons (Fig. 2). We found that the SLP and SIP received signals regarding the primary pheromone, whereas the LH received input mainly about the behavioral antagonist. The second pheromone was also represented in the LH, but with a rather low intensity. All female-produced substances were represented the VLP.

## Discussion

In moths, the sex pheromone is usually produced as a blend of several components in a species-specific ratio (Baker & Hansson, 2016; Christensen et al., 1995). Principally, the male is attracted by the major pheromone component released by a conspecific female. While minor components do not elicit upwind flight on their own; they may enhance attraction (Kehat & Dunkelblum, 1990), and can also serve as behavioral antagonists, since such components are often produced by heterospecific females (reviewed by Berg et al., 2014) or immature conspecific females (Chang et al., 2017). Despite these innate responses, the mate-searching activities, including an initial surge and zig-zag casting behavior (Cardé & Willis, 2008; Kuenen & Carde, 1994; Vickers & Baker, 1994), are not simple olfactory reflexes. The data presented here, comprising a large number of MGC medial-tract PNs, mainly originating from one of three easily identifiable glomeruli, indicated that such behavioral responses are related not only to spatial representation of odor valence in the lateral protocerebrum, but also to the intensity of the relevant signals.

### Pheromone signaling along parallel antennal-lobe tracts

In a recent study, we found that lateral-tract MGC PNs, which appear to originate from the cumulus only, convey a robust and rapid signal about the primary pheromone into a specific target area, i.e. the column in the SIP (Chu et al., 2020a). In the study presented here, we explored MGC-PNs in the two remaining tracts, the medial and the mediolateral. To clarify how the pheromone-information is carried along the three parallel tracts, we integrated the morphological findings from the previous study with our current results, and constructed a comprehensive map displaying the neural connections between MGC/AL glomeruli and the protocerebral target areas for each tract (Fig. 7A-B).

**Figure 7.**
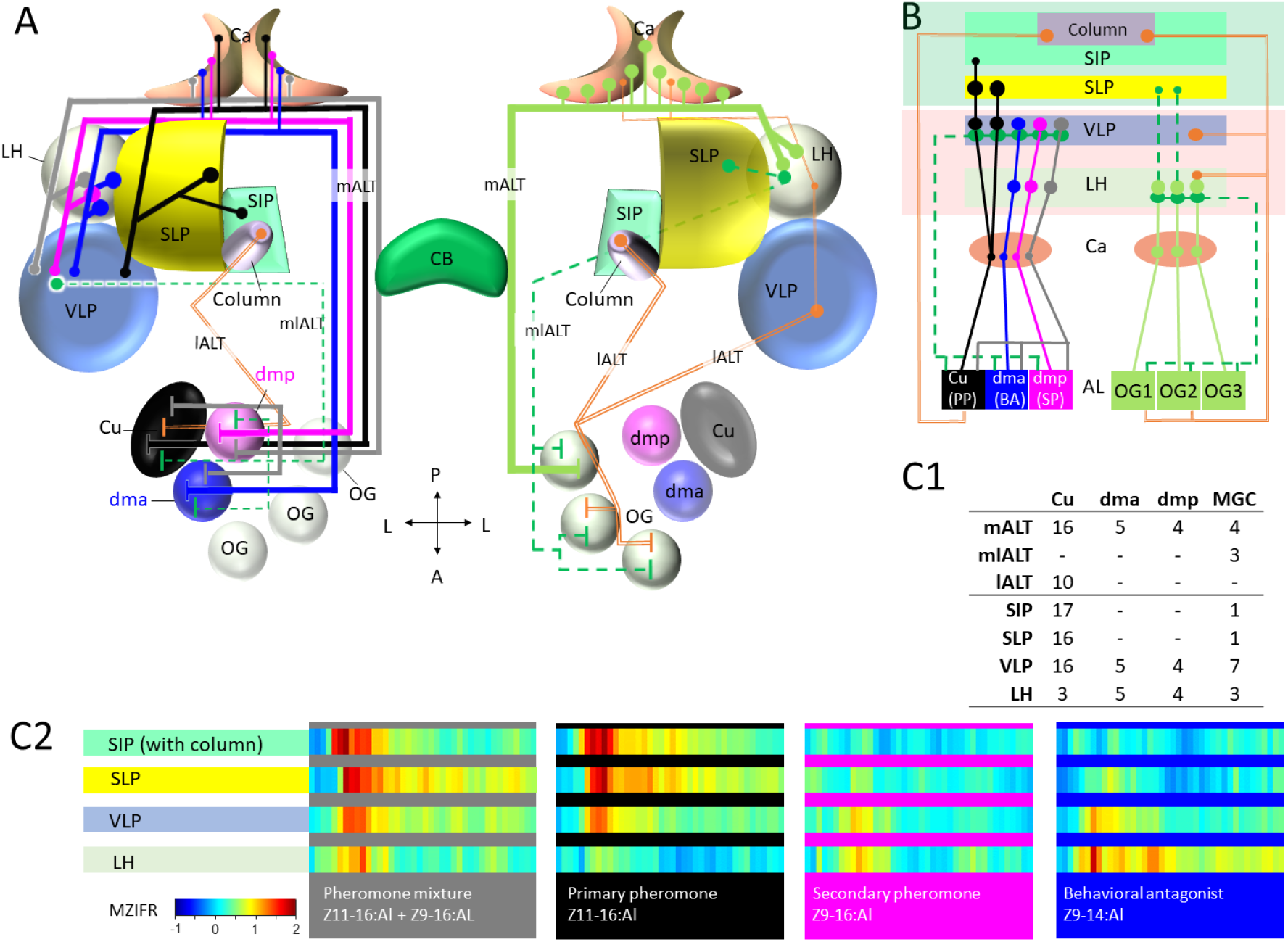
Summary of morphological and physiological features of MGC PNs across the three main tracts. (**A**) A graphic representation of the olfactory pathways in male moth brain. The morphological features of MGC PNs is displayed on the left hemisphere. Multiglomerular medial-tract MGC PNs (grey) and uniglomerular PNs arborizing in the dmp (magenta) or dma (blue) project to the calyces (Ca), anteroventral LH, and VLP, while the uniglomerular PNs with dendrites in the cumulus (Cu; black) target the Ca, VLP, SLP, and posterior SIP. The multiglomerular mediolateral-tract PNs (dashed green lines) target primarily the VLP, whereas the uniglomerular lateral-tract PNs innervating the cumulus runs directly to the column in the anterior SIP (Chu et al., 2020a). In the right hemisphere, a plethora of different PN-types arborizing in the ordinary glomeruli (OG) innervate several lateral protocerebral neuropils (Ian et al., 2016; Kymre et al., 2020). (**B)** Projection scheme of MGC PNs versus OG PNs in the higher brain regions. Three main antennal lobe tracts (ALTs) are illustrated: Solid line, medial ALT (mALT); Dash line, mediolateral ALT (mlALT); Double line, lateral ALT (lALT). (**C**) Summary of recorded and labelled PNs’ morphologies across the three main tracts, with the 10 lALT PNs being from our previous study (Chu et al., 2020a). All output neurons in (**C1)** were used to create an overview of temporal response properties in (**C2**), indicating mean responses (MZIFR; mean Z-scored instantaneous firing rate) to female-produced odorants. Note that these PNs were sampled from the antennal lobe, while the length of axons and action potential transmission rates have not been included, the actual timing of synaptic output onto the protocerebral neuropils is thereby not represented.

As pheromone-signaling appears to be particularly fast in lateral-tract MGC PNs (Chu et al., 2020a), we decided to illustrate how the temporal response properties of MGC PNs from the three main tracts are represented in the protocerebral neuropils. Interestingly, when including the data from lateral-tract MGC PNs (Chu et al., 2020a), we discovered a sequential and logic response pattern in the different protocerebral areas (Fig. 7B-C). Here, the excitatory response to the pheromone mixture arose first in the SIP, as a typical phasic response. After 20 ms, a phasic-tonic response to the binary mixture appeared in the SLP and VLP. Finally, about 120 ms later, a weak signal lasting for 10 ms could be seen in the LH. The primary pheromone evoked similar temporal response patterns in the SIP, SLP, and VLP, but barely any signal could be observed in the LH. The secondary pheromone elicited weak and delayed excitation in the VLP and LH, corresponding in time with the tiny peak evoked by the binary mixture in the LH. Finally, the behavioral antagonist induced a clear and long-lasting response in the LH and a considerably weaker activation in the VLP. Notably, the very fast response in the SIP region, which occurred during stimulation with the pheromone blend and the primary constituent exclusively involves the previously reported lateral-tract MGC-PNs terminating in the column (Chu et al., 2020a). Comparing the response onset in the SIP with the other output regions, demonstrates that the lateral-tract PNs react prior to PNs in the other main tracts (Fig. 7C).

### Protocerebral output-areas of MGC medial-tract neurons are organized according to MGC-unit origin

Despite the relatively wide-spread and partly overlapping terminal branches of the total assembly of medial-tract MGC-PNs, including 16 originating in the cumulus, 4 in the dmp, 4 in the dma, and 5 being multiglomerular, we discovered a pattern implying a spatial arrangement according to the cumulus-PNs versus the dma- and dmp-PNs. As discussed in detail below, this may correspond to the ‘lateral-medial’ representation of attraction- and aversion-related pheromone-signals previously reported in the protocerebrum of the closely related *Helicoverpa* species, *H. assulta*, and also in corresponding areas in the silk moth, *Bombyx mori* (Kanzaki et al., 2003; Seki et al., 2005; Zhao et al., 2014). In *H. armigera*, we found that signal representation related to attraction and inhibition is both segregated and integrated, but in distinct protocerebral neuropils.

#### Inputs related to sexual attraction are primarily processed in the SLP and SIP

All uniglomerular mALT PNs innervating the SLP and SIP had their dendrites in the cumulus. These cumulus-PNs responded mainly to the primary pheromone. Although the behavioral antagonist excited 19% (3 of 16) of the cumulus-PNs as well, these responses were not at all comparable to those associated with the primary pheromone. All data, including our calcium-imaging measurements (Fig. 1) and mean intracellular responses (Fig. 6A) confirm that the cumulus is devoted to process input about the primary pheromone. Notably, the multiglomerular mlALT PN innervating the SLP and SIP had its most dense innervations in the cumulus as well (Fig. 2L). Thus, third-order pheromone-processing neurons in the SLP and SIP should primarily be involved in computations related to attraction.

#### Anteroventral parts of the LH process dma and dmp input

In contrast to the SLP and SIP, being innervated by the medial-tract cumulus-neurons, the anteroventral LH was the target for the PNs originating in one of the two smaller MGC-units, the dma and dmp. Thereby, input about the primary pheromone is processed dorsomedially to input about the secondary component and the behavioral antagonist. This lateral-medial separation resembles the spatial pattern previously reported in the closely related species, *H. assulta*, where the primary pheromone is represented medially to the interspecific signal (Zhao et al., 2014). In the “older” species *H. assulta*, however, the secondary pheromone is not processed together with the behavioral antagonist, but with the primary pheromone.

Summarizing the response patterns of the MGC PNs innervating the LH (Fig. 6B & 7C2), we found that the highest mean firing rate appeared during application of the behavioral antagonist. The secondary pheromone (plus the pheromone blend) also induced increased spike frequency, but substantially weaker. As mentioned in the introduction, the behavioral antagonist, Z9-14:Al, reduces mating-associated behaviors at high concentrations (Kehat & Dunkelblum, 1990; Wu et al., 2015), while it enhances such behaviors at low concentrations (Wu et al., 2015; Zhang et al., 2012). Indeed, low concentrations of this component is released by *H. armigera* females (Kehat & Dunkelblum, 1990; Zhang et al., 2012), whilst sympatric *Heliothis peltigera* females emit higher concentrations of the signal (Hillier & Baker, 2016). The secondary pheromone component, Z9-16:Al, resembles Z9-14:Al in that it may also include both enhancement and inhibition of attraction, dependent on the concentration. Concretely, Z9-16:Al is known to augment attraction when combined with the primary pheromone, but to reduce attraction when constituting more than 10 % of the binary pheromone mixture (Kehat & Dunkelblum, 1990; Kehat et al., 1980). Notably, high concentrations of Z9-16:Al may serve as an interspecific signal, due to it being the primary pheromone of sympatric *H. assulta* females (Cork et al., 1992). Altogether, in light of the complex behavioral functions of Z9-16:Al and Z9-14:Al, our data indicate that the anteroventral part of the LH is involved in inhibition of attraction when the moth encounters high concentrations of the relevant components.

#### A proposed interaction between the LH and the SLP/SIP

How can each of the two substances, Z9-14:Al and Z9-16:Al, serve opposite functions? During mate-searching behavior, the likelihood of encountering the primary pheromone is much larger than detecting the minor components, due to their proportion ratios in the natural female-released pheromone mixture. As such, the SLP and SIP, which receive information about the primary pheromone, is likely to be activated prior to the anteroventral LH. Notably, a prominent neural link between the LH and SLP in moths (Namiki & Kanzaki, 2019) appears to be a strong candidate for governing the “correct” mate-searching behavior. A study in the fruit fly has demonstrated that the LH output projects primarily to SLP and secondly to the SIP, and about one third of these neurons are GABAergic/glutamatergic (Dolan et al., 2019). Since both neurotransmitters are inhibitory (Liu & Wilson, 2013), this indicates that LH input may block the activation of SLP and SIP output neurons and their downstream circuits. The interaction between the LH and SLP/SIP could act as a Boolean logic gate implementing the AND-NOT function, only passing information downstream when aversive signaling is minimally present in the anteroventral LH.

We propose that this form of Boolean logic represents a flexible interaction between the LH and SLP/SIP. The LH output is mainly modulated by the input from the AL PNs, and the activity of these neurons increases with increasing odor concentration (Gupta & Stopfer, 2012; Lerner et al., 2020; Sachse & Galizia, 2003). When a moth encounters the primary pheromone and a very low concentration of the secondary pheromone (released by conspecific female), the LH input is weak and the inhibition from the LH to SLP/SIP minimal, therefore, the output from the SLP/SIP to downstream targets remains strong. Should a big amount of the secondary pheromone or the behavioral antagonist (released by a heterospecific female) recruit vigorous MGC PN output onto the LH, the GABAergic/glutamatergic LH neurons will inhibit the activity in SLP/SIP and the downstream signaling. This might be perceived as a loss of contact with the primary pheromone, leading the moth to quickly engage in a casting flight, i.e. crosswind flight with no net upwind movement (Kuenen & Carde, 1994), and thereby promote an appropriate mate-searching strategy.

Given that the SLP/SIP is involved in processing pheromone information solely about attraction, while the LH represents concentration-dependent ambiguous signals, one question appearing is how the presence of low-concentrations of the minor components enhances attraction when added to the primary pheromone. Concerning Z9-16:Al, for example, our findings indicate that integration of the secondary and primary pheromone might occur at the level of MGC output PNs. Here, we first analyzed the physiological data of 11 uniglomerular medial-tract PNs with projections to SLP/SIP having excitatory responses to the primary pheromone and pheromone mixture. Their response amplitudes as well as response peaks were compared during stimulation with the primary pheromone alone and the binary mixture (secondary pheromone < 5%). In both electrophysiological recording and calcium-imaging experiments, the pheromone mixture showed a tendency of evoking a stronger response (Fig. 13E). We also found at least three cumulus-PNs with projections in the SLP that had a tonic and delayed excitation to the secondary pheromone (Fig. 4 and Fig. 12).

#### The pheromone-processing role of the VLP is complex

Unlike the neuropils discussed above, the VLP was innervated by all labelled MGC PNs. Specifically, they sent terminal projections into a compact sub-domain of the dorsoanterior VLP, which thereby gets intermingled signals about all female-relevant compounds. This suggests that the VLP is involved in combinatorial coding, possibly recognizing the optimal species-specific signal. In addition, combinatorial coding of pheromone signals seems to occur at an earlier level as well. The dendritic organization of the multiglomerular MGC PNs prime these neurons for combinatorial coding. This was, perhaps, most clearly demonstrated by the broad response profiles of the mediolateral-tract PNs with evenly distributed dendrites across the MGC (Fig. 4 and Fig. 4 - figure supplement 7). These PNs responded to all female-produced components, but with distinct temporal patterns (Fig. 5), and may provide excitatory or inhibitory input to the medial-tract PNs in the VLP since about half of the axons forming this tract is GABAergic (Berg et al., 2009). Furthermore, we demonstrated that these mlALT PNs had overlapping terminals with the medial-tract MGC PNs in this neuropil (Fig. 3A). As the VLP is the only known region with overlapping terminals of pheromone PNs across different tracts, this makes it a particularly interesting neuropil for future studies investigating parallel processing in the pheromone system.

## Materials and methods

### Insects

Male moths (2-3 days) of *H. armigera* were used in this study. Pupae were purchased from Keyun Bio-pesticides (Henan, China). After emergence, the moths were kept at 24 °C and 70% humidity on a 14:10 h light/dark cycle, with 10% sucrose solution available ad libitum. According to Norwegian law of animal welfare, there are no restrictions regarding experimental use of Lepidoptera.

### Calcium imaging

Totally, 8 males (2-3 days) were used in calcium-imaging experiments. Retrograde selective staining of MGC PNs has been reported elsewhere (Chu et al., 2020a; Ian et al., 2017; Sachse & Galizia, 2002; Sachse & Galizia, 2003). A glass electrode tip coated with Fura-2 dextran (potassium salt, 10,000 MW, Molecular Probes) was inserted into the calyces to selectively label the medial-tract PNs. The insects were then kept for 12 h at 4 °C before the experiment, to facilitate retrograde transportation.

In vivo calcium imaging recordings were obtained with an epifluorescent microscope (Olympus BX51WI) equipped with a 20x/1.00 water immersion objective (OlympusXLUMPlanFLN). Images were acquired by a 1344×1224 pixel CMOS camera (Hamamatsu ORCA-Flash4.0 V2 C11440-22CU). The preparation was excited with 340 nm and 380 nm monochromatic light, respectively (TILL Photonics Polychrome V), and data were acquired ratiometrically. A dichroic mirror (420nm) and an emission filter (490-530nm) were used to separate the excitation and emission light. Each recording consisted of 100 double frames at a sampling frequency of 10 Hz with 35 ms and 10 ms exposure times for the 340 nm and 380 nm lights, respectively. The duration of one recording trial was 10 s, including 4 s with spontaneous activity, 2 s odor stimulation, and a 4 s post-stimulus period. The odor stimulation was carried out by a stimulus controller (SYNTECH CS-55), via which humidified charcoal filtered air was delivered through a 150 mm glass Pasteur-pipette with a piece of filter paper containing the stimulus. Each odor stimulus was applied twice. The interval between trials was 60 s to avoid possible adaptation.

### Intracellular recording and staining

Preparation of the insect has been described in detail elsewhere (KC et al., 2020; Kymre et al., 2020). Briefly, the moth was restrained inside a plastic tube with the head exposed and then immobilized with dental wax (Kerr Corporation, Romulus, MI, USA). The brain was exposed by opening the head capsule and removing the muscle tissue. The exposed brain was continuously supplied with Ringer’s solution (in mM): 150 NaCl, 3 CaCl2, 3 KCl, 25 sucrose, and 10 N-tris (hydroxymethyl)-methyl-2-amino-ethanesulfonic acid, pH 6.9).

The procedure of intracellular recording/staining of neurons was performed as previously described (Chu et al., 2020a; Ian et al., 2016; Zhao et al., 2014). Sharp glass electrodes were made by pulling borosilicate glass capillaries (OD: 1 mm, ID: 0.5 mm, with hollow filament 0.13 mm; Hilgenberg GmbH, Germany) on a horizontal puller (P97; Sutter Instruments, Novarto, CA, USA). The tip of the micro-pipette was filled with a fluorescent dye, i.e. 4% biotinylated dextran-conjugated tetramethylrhodamine (3000 mw, micro-ruby, Molecular Probes; Invitrogen, Eugene, OR, USA) in 0.2 M KAc. The glass capillary was back-filled with 0.2 M KAc. A chlorinated silver wire inserted into the muscle in the mouthpart served as the reference electrode. The recording electrode, having a resistance of 70–150 MΩ, was carefully inserted into the dorsolateral region of the AL via a micromanipulator (Leica). Neuronal spike activity was amplified (AxoClamp 1A, Axon Instruments, Union, CA, USA) and monitored continuously by oscilloscope and loudspeaker. Spike2 6.02 (Cambridge Electronic Design, Cambridge, England) was used as acquisition software. During recording, the moth was ventilated constantly with a steady stream of fresh air. During odor stimulation, a pulse of air from the continuous airstream was diverted via a solenoid-activated valve (General Valve Corp., Fairfield, NJ, USA) through a glass cartridge bearing the odorant on a piece of filter paper. Up to six odors were tested in each recording experiment, while the number of trials were dependent on neuronal contact. The stimulation period was 400 ms. After testing all odor stimuli, the neuron was iontophoretically stained by applying 2–3 nA pulses with 200 ms duration at 1 Hz for about 5–10 min. In order to allow neuronal transportation of the dye, the preparation was kept overnight at 4 °C. The brain was then dissected from the head capsule and fixed in 4% paraformaldehyde for 1-2 h at room temperature, before it was dehydrated in an ascending ethanol series (50%, 70%, 90%, 96%, 2 × 100%; 10 min each). Finally, the brain was cleared and mounted in methylsalicylate.

### Odor stimulation

During intracellular recordings, the following stimuli were tested: (i) the primary sex pheromone of *H. armigera*, Z11-16:Al, (ii) the secondary sex pheromone, Z9-16:Al, (iii) the binary mixture of Z11-16:Al and Z9-16:Al, (iv) the behavioral antagonist of *H. armigera*, Z9-14:Al, (v) the head space of a host plant (sunflower leaves), and (vi) hexane as a vehicle control. The three insect-produced components were obtained from Pherobank (Wijk bij Duurstede, Netherlands). Stimuli i-iv were used in two distinct stimulations protocols, i.e. either with low or high concentrations of the respective stimuli. In the low concentration protocol, the mixture of Z11-16:Al and Z9-16:Al were in a 95:5 proportion, while the high concentration protocol used a 97:3 proportion, both resembling the natural blend emitted by conspecific females (Hillier & Baker, 2016; Kehat et al., 1980; Piccardi et al., 1977; Wu et al., 1997). The insect-produced stimuli were diluted in hexane (99%, Sigma), and applied to a filter paper that was placed inside a 120 mm glass cartridge. For the low concentration protocol, the final amount per filter paper was 10 ng of the relevant stimulus, whereas the high concentration protocol included 100 ng. Note that the ID of PNs stimulated with the low concentration protocol were numbered (e.g. Cu-mALT**1**), while the PNs stimulated with the high stimulation protocol were named with letters (e.g. Cu-mALT**a**). To avoid adaptation, PNs in the high concentration protocol did not receive repeated trials, as large dosages of pheromones are associated with prolonged adaptation of the relevant OSNs (Dolzer et al., 2003).

The same odor stimuli as listed above were used during the calcium imaging experiment, but at a higher concentration (required to evoke a response in this technique), i.e. 10 μg at the filter paper. To avoid adaptation, the interstimulus interval was 1 min. An additional stimulus containing a 50:50 mixture of host plant (20μl) and pheromone mix (20μl 95:5 mixture) was added in the calcium imaging measurements.

### Confocal microscopy

Whole brains were imaged by using a confocal laser-scanning microscope (LSM 800 Zeiss, Jena, Germany) equipped with a Plan-Neofluar 20x/0.5 objective and/or a 10x/0.45 water objective (C-achroplan). Micro-ruby staining was excited with a HeNe laser at 553 nm and the fluorescent emission passed through a 560 nm long-pass filter. In addition to the fluorescent dyes, the auto-fluorescence of endogenous fluorophores in the neural tissue was imaged to visualize relevant structures in the brain containing the stained neurons. Since many fluorophore molecules were excited at 493 nm, auto-fluorescent images were obtained by using the argon laser in combination with a 505-550 nm band pass filter. Serial optical sections with resolution of 1024 x 1024 pixels were obtained at 2-8 μm intervals. Confocal images were edited in ZEN 2 (blue edition, Carl Zeiss Microscopy GmbH, Jana, Germany) and MATLAB R2018b.

### Reconstruction of individual neurons and nomenclature

For visualization purposes, each successfully stained neuron was reconstructed in AMIRA 5.3 (Visualization Science Group) by using the *SkeletonTree* plugin (Evers et al., 2005; Schmitt et al., 2004) and by this the morphology of the neuronal filaments and their thickness were digitally reproduced. Further, the reconstructed neurons were manually transformed into the 3D representative brain, as used previously (Chu et al., 2020a; Chu et al., 2020b), for illustration. Based on studies in the silk moth, *B. mori*, the “delta region of the inferior lateral protocerebrum” (ΔILPC), a pyramid-shaped area formed by medial and mediolateral tract PNs, was previously described as a main MGC-output area (e.g. Seki et al., 2005). The current standard insect brain nomenclature, established by Ito et al. (2014), does not include the ΔILPC, which seems to be part of several protocerebral neuropils (Ito et al., 2014; Lee et al., 2019; Lei et al., 2013). The identification and naming of neuropil structures in the representative brain has been adapted from the standard nomenclature established by Ito et al. (2014). The orientation of all brain structures is indicated relative to the body axis of the insect, as in Homberg et al. (1988).

### Calcium imaging data analysis

In this study, responses of medial-tract PNs innervating each MGC unit were analyzed. Recordings were acquired with Live Acquisition V2.3.0.18 (TILL Photonics GmbH, Kaufbeuren, Germany) and imported in KNIME Analytics Platform 2.12.2 (KNIME GmbH, Konstanz, Germany). Here, ImageBee neuro plugin (Strauch et al., 2013) was used to construct AL maps and glomerular time series. To determine an average baseline activity, the Fura signal representing the ratio between 340 and 380 nm excitation light (F340/F380) from 0.5 to 2.5 s (frames 5-25, within 4 s spontaneous activity) was selected and set to zero. Neuronal activity traces were thus represented as changes in fluorescent level, specified as ΔF_340_/F_380_. We defined a *Threshold* based upon the control (hexane) traces across 8 individuals at a 5% significance level: 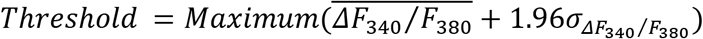. Responses were defined when the mean peak of ΔF_340_/F_380_ trace (n=8) for distinct stimuli was higher than *Threshold*. For displaying the response amplitude for each stimulus, the averaged ΔF340/F380 within 2s stimulation window was used.

### Spike data analysis

The electrophysiological data was spike-sorted and analyzed in Spike 2.8. Each odor application trial comprised a total period of 2.4 s, including a 1 s pre-stimulation window for baseline activity prior to the stimulus onset, 0.4 s stimulation period, and 1 s post-stimulation period. For describing the temporal neural activity, the Z-scored instantaneous firing rates (ZIFR) of every 10 ms for each trial were registered. To measure stimulus-specific responses of individual PNs, the odor-evoked response properties were analyzed in the mean ZIFR (MZIFR) across repetitive trials with the same stimulus. Significant responses were determined by the upper threshold (TU) and lower threshold (TL) calculated according to the mean MZIFR in the 1 s pre-stimulation window (MZIFR_PS_) at a 5% significance level:

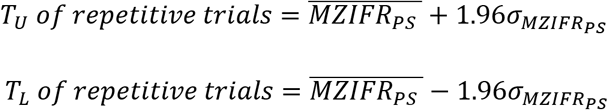

If there was an individual ZIFR in the stimulation window higher than the value of T_U_ or lower than T_L_, the stimulation was determined as evoking excitatory or inhibitory response, respectively.

To illustrate the response patterns, the stimulation window was divided into two sub-windows (SW), the SW(i) included the first 100 ms of the stimulus application and SW(ii) included the remaining 300 ms of the stimulus application. If one of the mean ZIFR during the stimulation sub-windows was higher than the value of ‘Upper Threshold’ or lower than ‘Lower Threshold’, the change in activation was determined as an excitatory or inhibitory response, respectively.

For displaying the response amplitude for each stimulus, we first standardized the baseline activity by setting the MZIFR before stimulation onset to zero. The response amplitude was then quantified as the ΔMZIFR averaged within two stimulation sub-windows, respectively. The ΔMZIFR in a 200ms post-stimulation window was also computed. For each trial, the onset of an excitatory response was determined at the time point when the Z-scored instantaneous firing rate (binned every 1ms) exceeded the corresponding response threshold (same formula as TU). The mean responses of PNs within groups categorized by the dendritic or axonal innervations were also computed.

## Acknowledgments

We thank Stanley Heinze for his contribution to the representative brain atlas, Baiwei Ma for assistance with data collection, and Tom Knudsen for technical support.

## Figure supplements

**Figure 1 – figure supplement 1.**
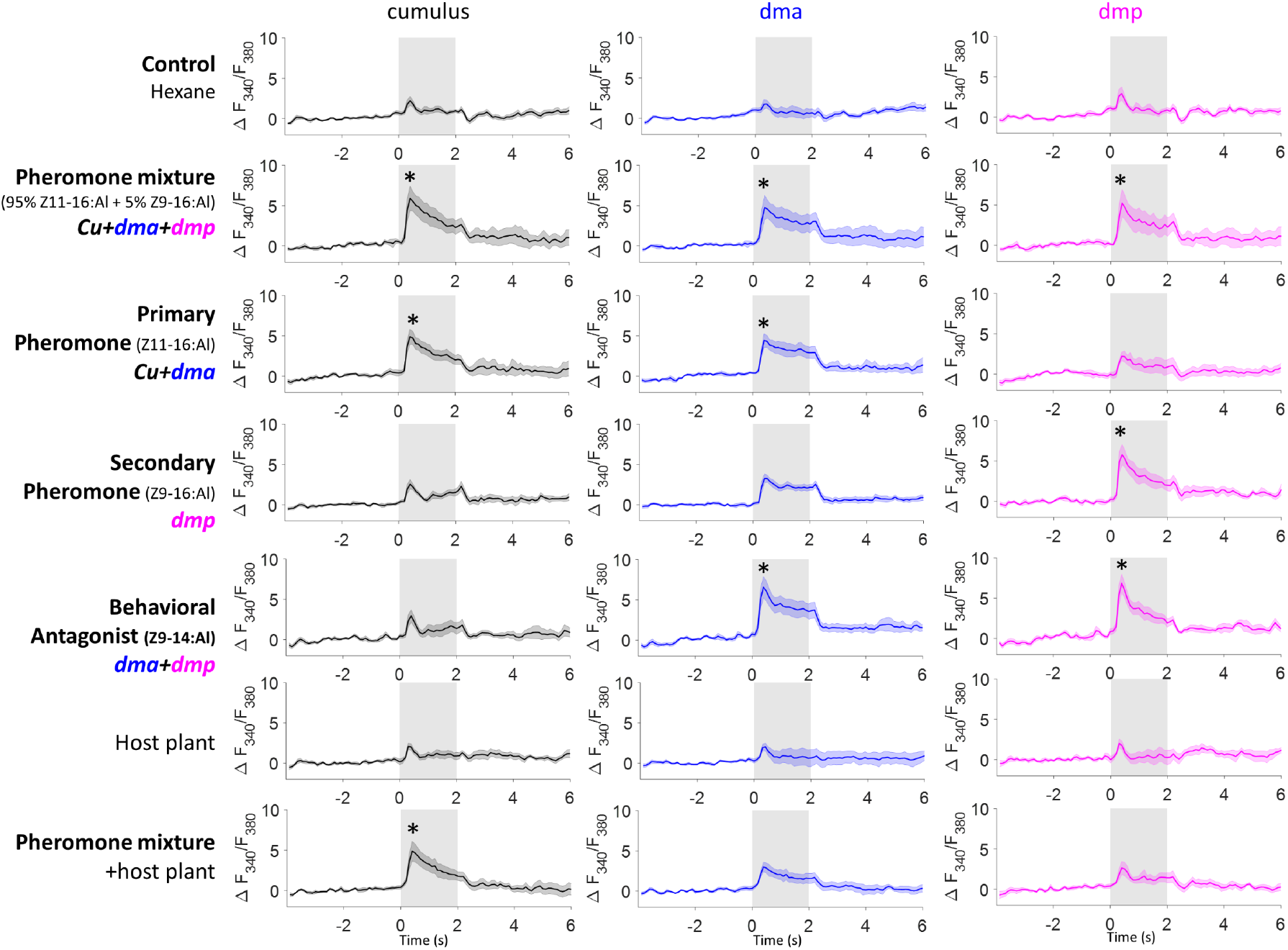
Mean response traces of the MGC units during stimulation with all presented stimuli, where * represents a statistically significant deviation from the Fura signal evoked by control.

**Figure 1 – figure supplement 2.**
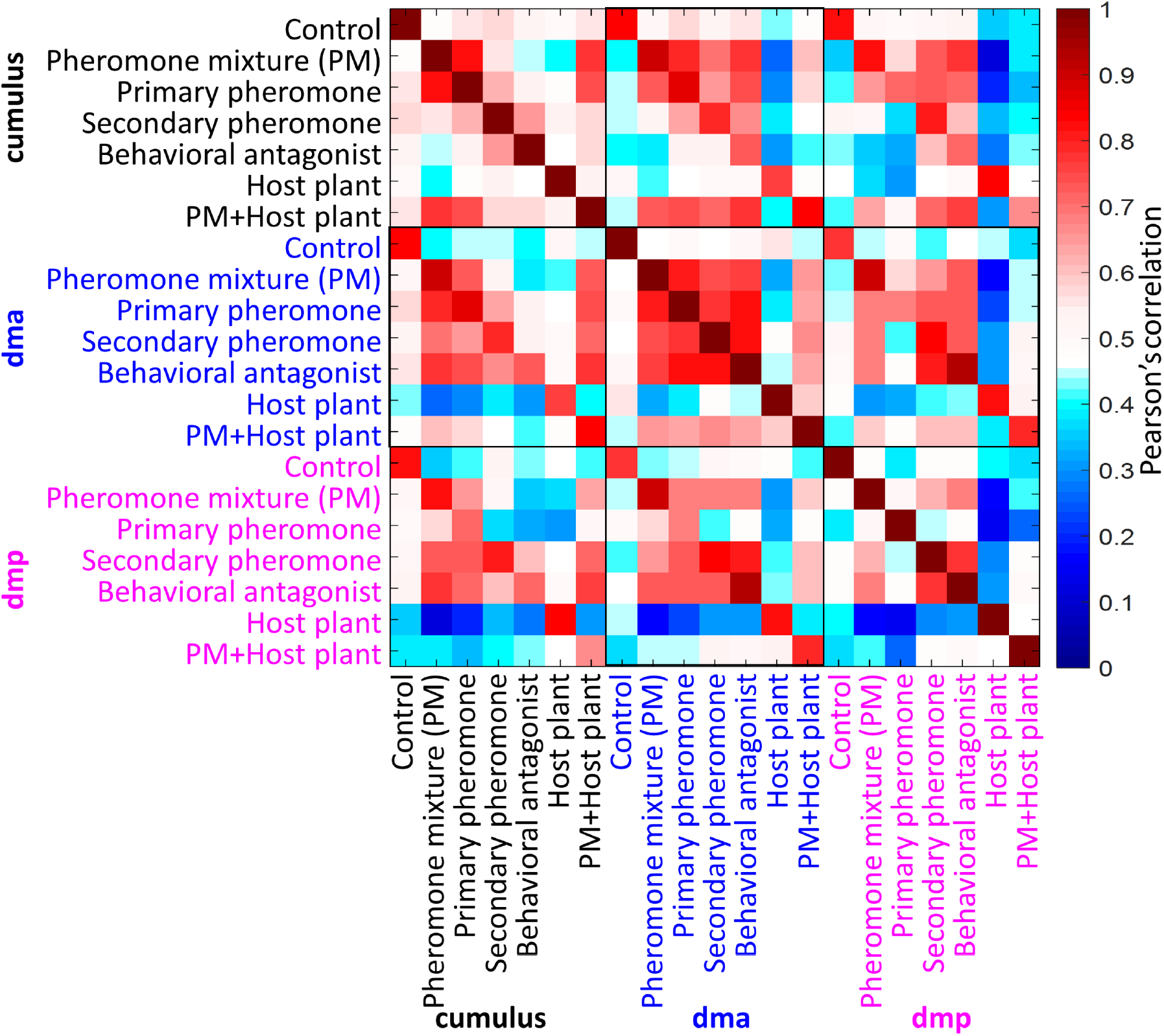
Cross-stimuli correlation plot of mean response traces of the MGC units during stimulation with all presented stimuli.

**Figure 2 – figure supplement 1.**
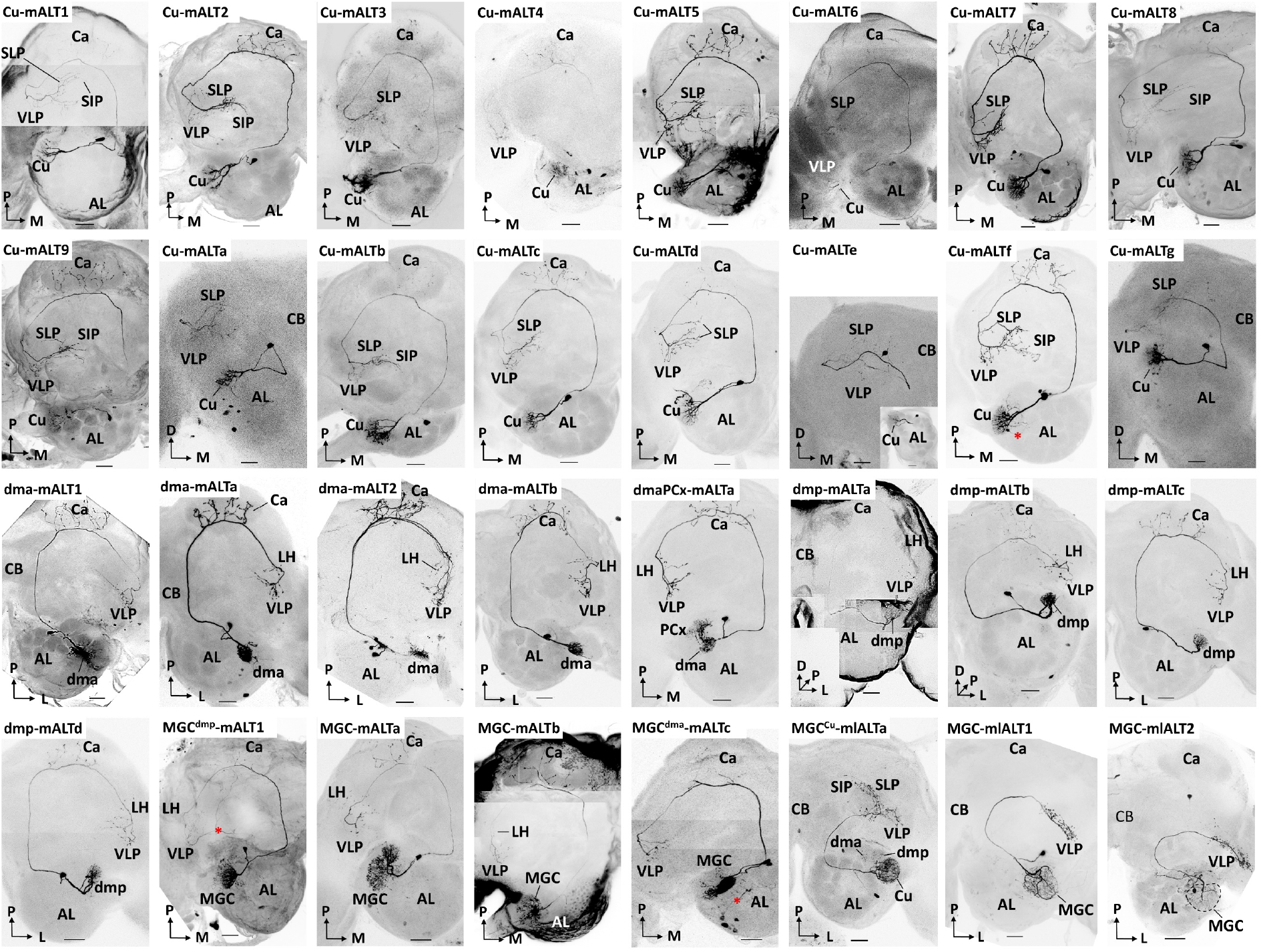
Morphology of all individually recorded PNs. Red asterisks indicate weakly co-labelled neurons. Scale bars: 50μm

**Figure 4 – figure supplement 1.**
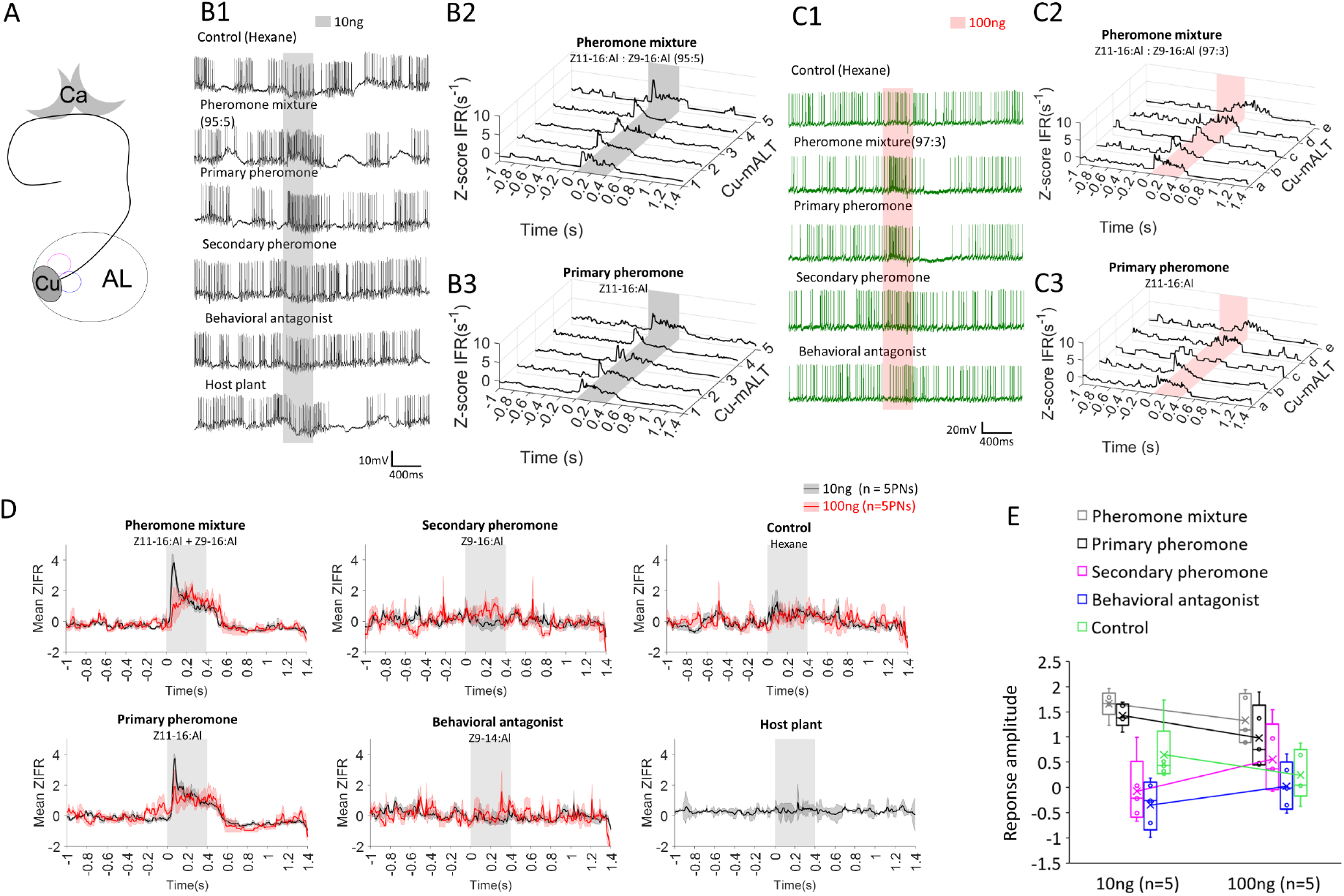
The predominant response pattern of mALT PNs originating in the cumulus. (**A**) Simplified scheme of a mALT PN innervating the cumulus. (**B1**) Spiking activity of one of the mALT cumulus neurons during application of odor stimuli at low concentration (10 ng). (**B2-B3**) Mean traces of Z-scored instantaneous firing rates (ZIFR) across repeated trials to the low concentration pheromone mixture (**B2**) and the primary pheromone (**B3**) in 5 cumulus neurons, Cu-mALT1-5. (**C1**) Spiking activity of one of the mALT cumulus neurons during application of odor stimuli at high concentration (100 ng). (**B2-B3**) Single trial traces of Z-scored instantaneous firing rates (ZIFR) to the high concentration pheromone mixture (**C2**) and the primary pheromone (C**3**) in 5 cumulus neurons, Cu-mALTa-e. (**D**) Comparison of mean ZIFR traces between low and high concentration protocols. (**E**) Box plots of response amplitude (ΔMZIFR) of the entire 400ms stimulation window in two concentration protocols. The response amplitudes across stimuli appeared more pronounced in the low concentration protocol, suggesting that impaired odor discrimination may occur at high odor concentrations.

**Figure 4 – figure supplement 2.**
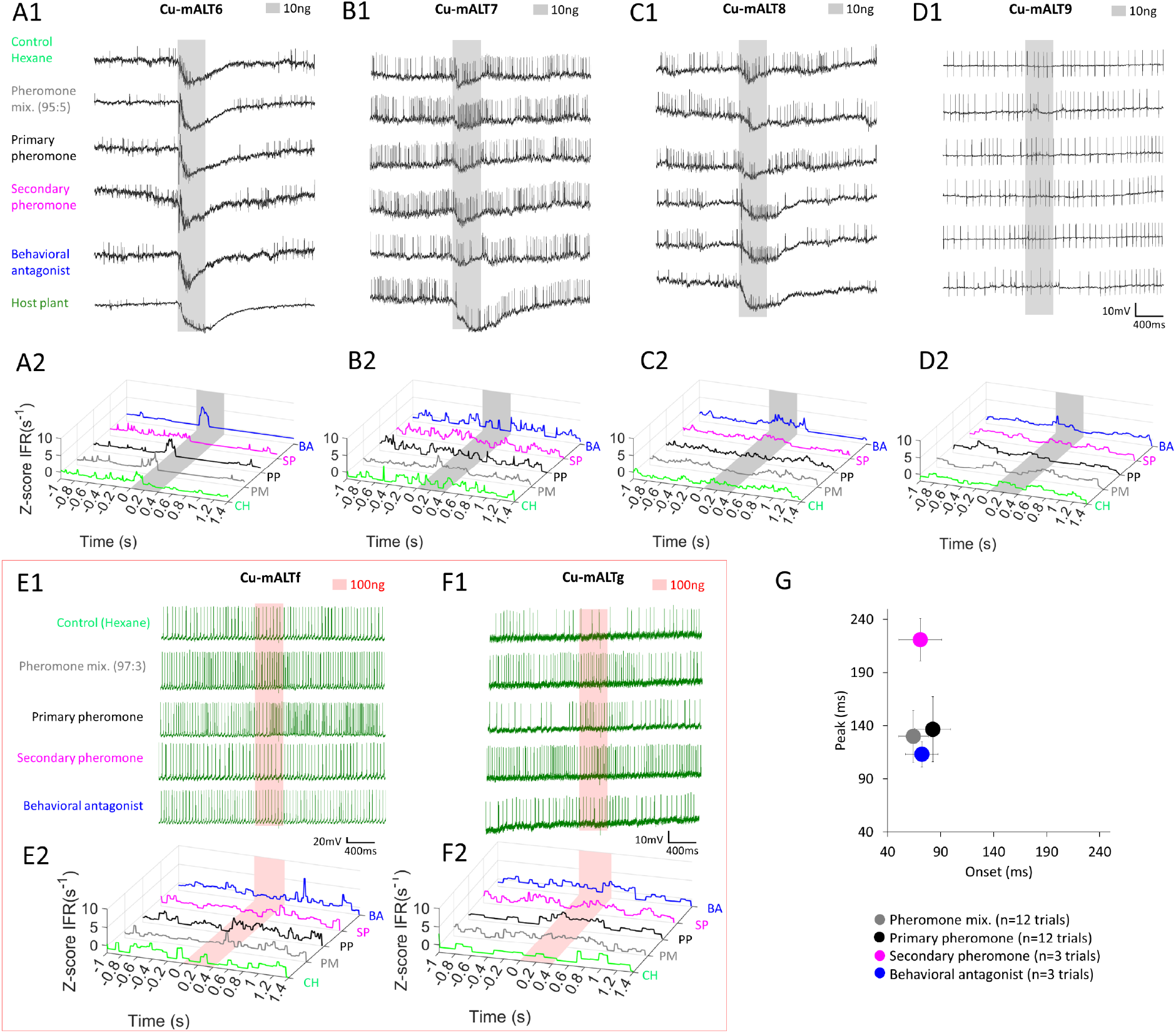
Heterogeneous response profiles of cumulus-innervating mALT PNs with homogenous morphologies. (**A-D**) Spiking activity and traces of instantaneous firing rates (ZIFR) in PNs with ID CU-mALT6-9 during application of odor stimuli at low concentration (10 ng). (**E-F**) Spiking activity and traces of instantaneous firing rates (ZIFR) in PNs CU-mALTf-g during application of odor stimuli at high concentration (100 ng). (**G**) Onset-Peak scatter plot including latency data from individual trials across the 6 presented PNs in (**A-F**). Data is presented as mean±se. The delayed peak was only evoked by the secondary pheromone. CH, hexane; PM, pheromone mixture; PP, primary pheromone; SP, secondary pheromone; and BA, behavioral antagonist.

**Figure 4 – figure supplement 3.**
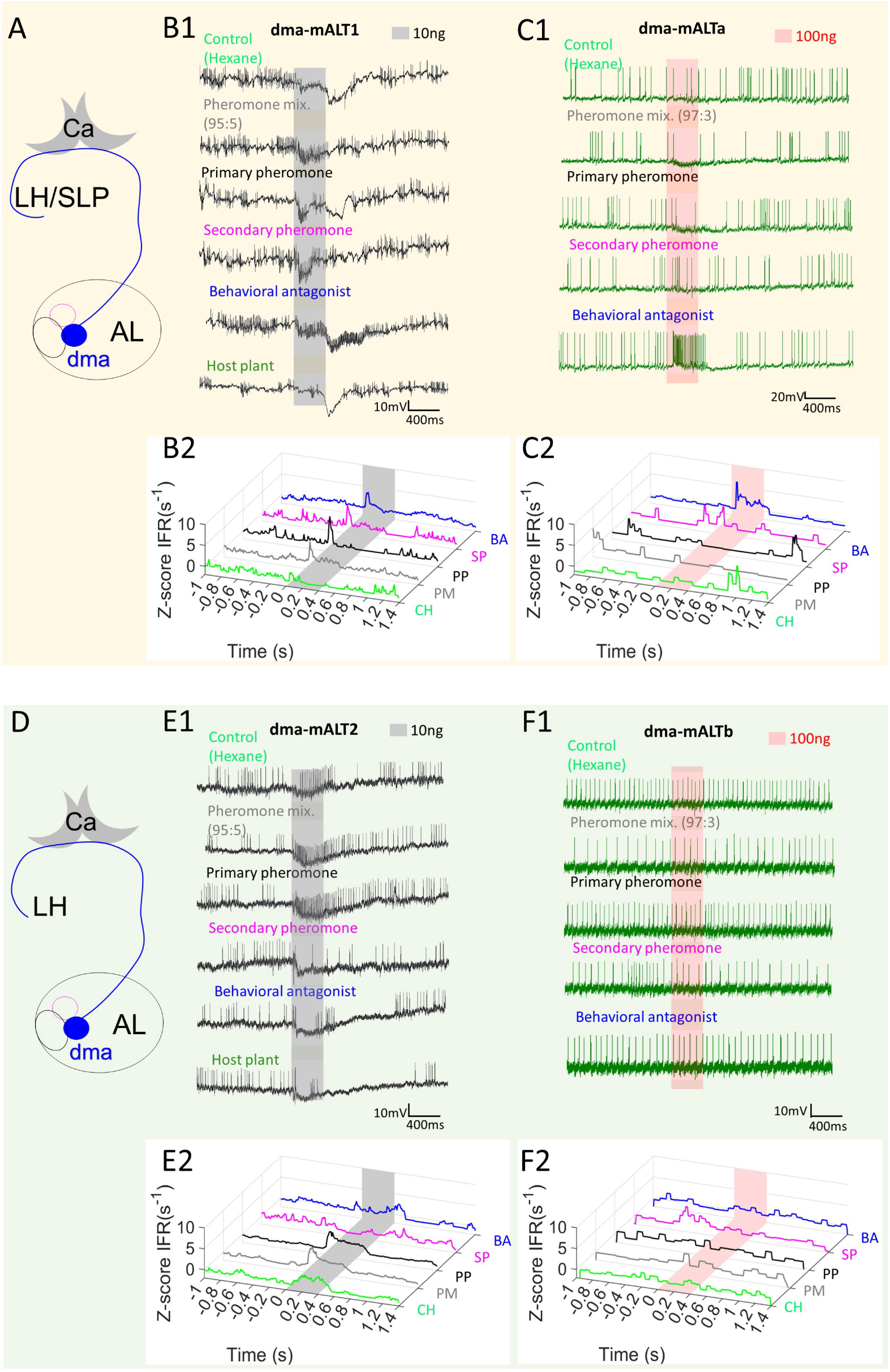
Heterogeneous response profiles of two types of dma-innervating mALT PNs. (**A-C**) Physiological profile of dma-PN type with extended terminals approaching to the SLP. The spiking activity and Z-scored instantaneous firing rates (ZIFR) trace showed that both PNs of this type were excited by the behavioral antagonist (BA) and the secondary pheromone (SP). PN dma-mALT1 in (**B**) was widely tuned, responding with phasic excitation to all stimuli except the control hexane (CH). PN dma-mALTa appeared to be suppressed by the pheromone mixture (PM) and the primary pheromone (PP). (**D-F**) Physiological profile of the other dma-PN type with restricted terminal regions. The spiking activity and Z-scored instantaneous firing rates (ZIFR) trace of the two relevant neurons illustrated that neither of them responded to the BA at low concentration (**E**) or high concentration (**F**).

**Figure 4 – figure supplement 4.**
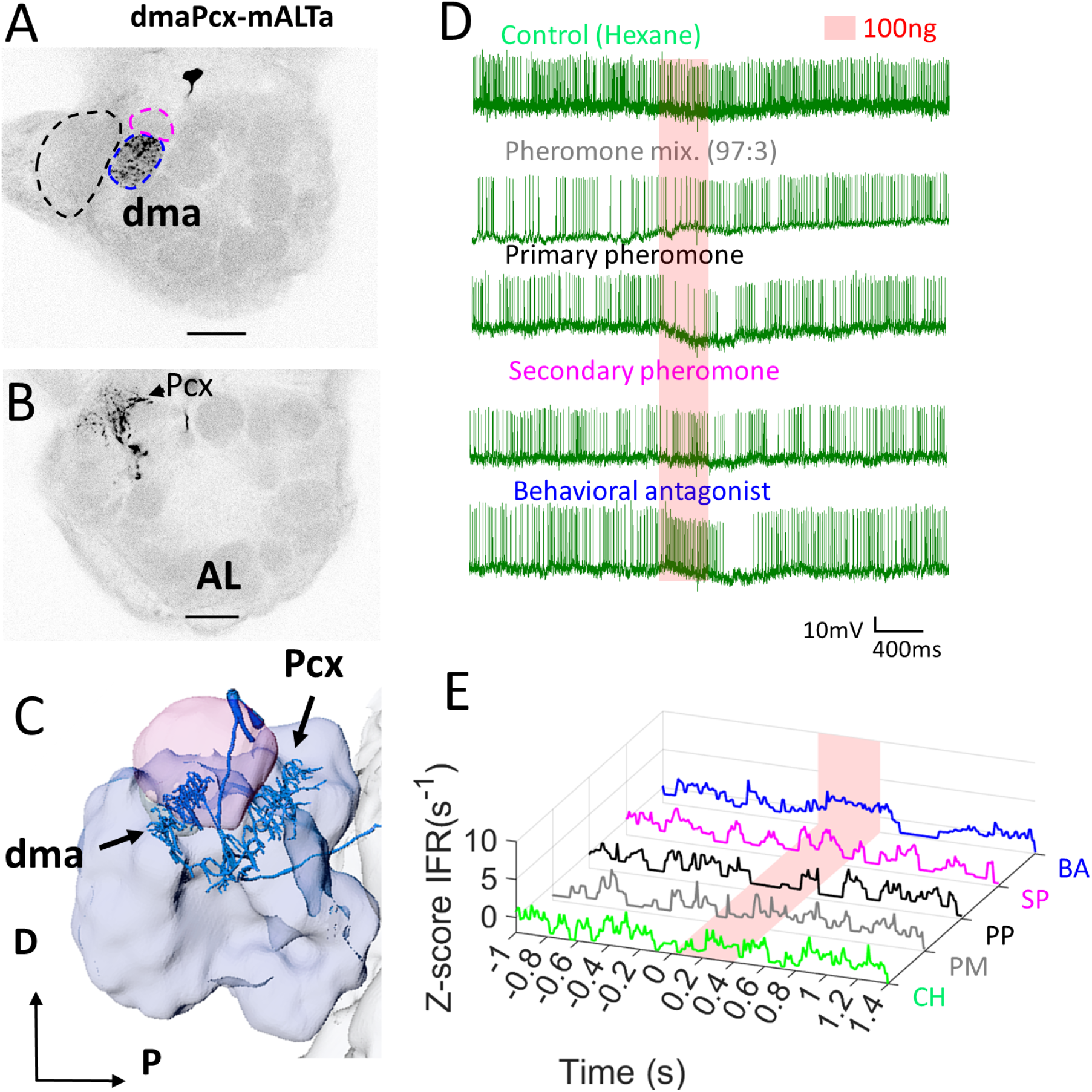
Dendritic morphology and response profile of the dmaPCx-mALT PN. (**A-B**) Confocal images demonstrate that this PN innervated the dma (blue dashed circle in **A**), but not the cumulus or dmp (black and pink dashed circles, respectively). Three posterior complex (PCx) glomeruli also received some dendritic processes (**B**). (**C**) Reconstruction demonstrating the AL innervation in sagittal view). (**D-E**) Spiking activity and Z-scored instantaneous firing rates (ZIFR) trace showed no responses that were significantly different from baseline firing rates. However, the firing rate of this PN was slightly enhanced when stimulated with the behavioral antagonist (BA), and moderately inhibited by the primary pheromone (PP). Scale-bars = 50 μm.

**Figure 4 – figure supplement 5.**
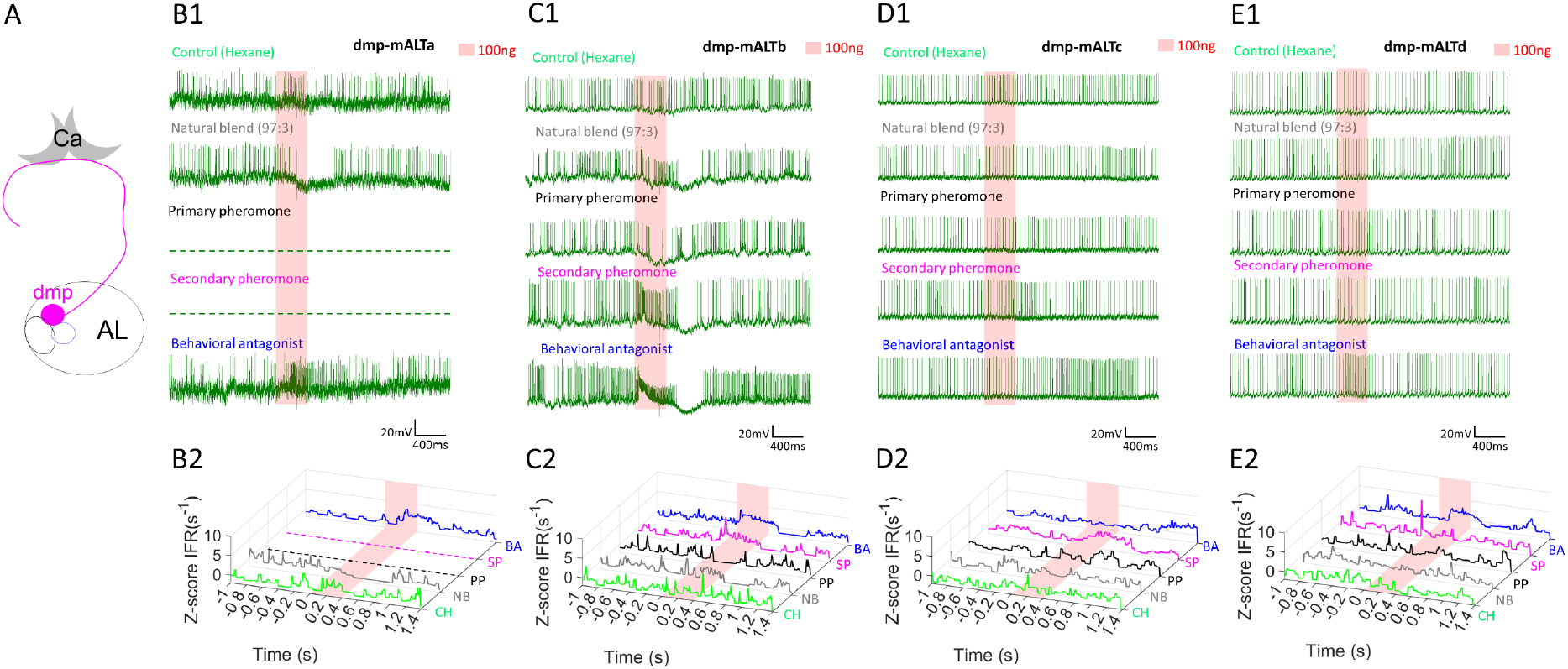
Response profiles of mALT PNs with uniglomerular dendrites in the dmp. (**A**) Schematic display of a mALT PN innervating the dmp. (**B-E**) Spike traces and standardized instantaneous firing rates (ZIFR) during stimulation with high concentrations of the female-produced stimuli indicated that the most prominent excitatory responses were elicited by the behavioral antagonist (BA) and/or the secondary pheromone (SP).

**Figure 4 – figure supplement 6.**
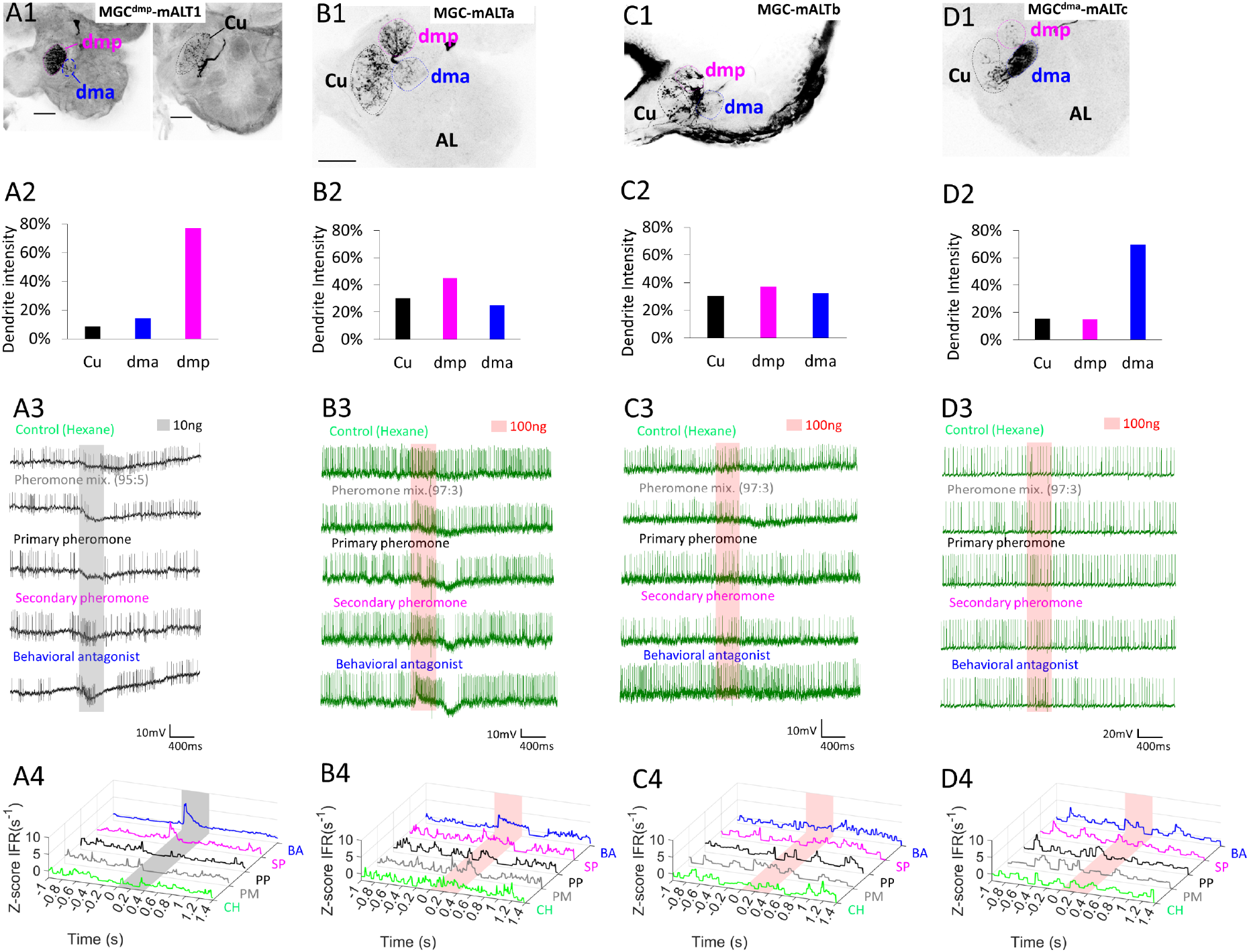
Dendritic MGC innervation and physiological features of multiglomerular mALT PNs. (**A**) The PN MGCdmp-mALT1 had the majority of its dendrites in the dmp, along with minor innervations in the dma and cumulus (Cu; **A1-A2**). This neuron responded with phasic excitations to both the secondary pheromone (SP) and the behavioral antagonist (BA), and was inhibited by the pheromone mixture (PM) and the primary pheromone (PP; **A3-A4**). (**B**) The dendrites of PN MGC-mALTa was quite evenly distributed across the MGC, yet this PN was strongly excited only by the BA stimulus. (**C**) PN MGC-mALTb also had evenly distributed dendrites, but had no strong responses to any of the presented stimuli. (**D**) The dendrites of PN MGCdma-mALTc innervated the dma densely, while the cumulus and dmp were sparsely innervated. This PN displayed no changes in firing rates upon odor stimulation.

**Figure 4 – figure supplement 7.**
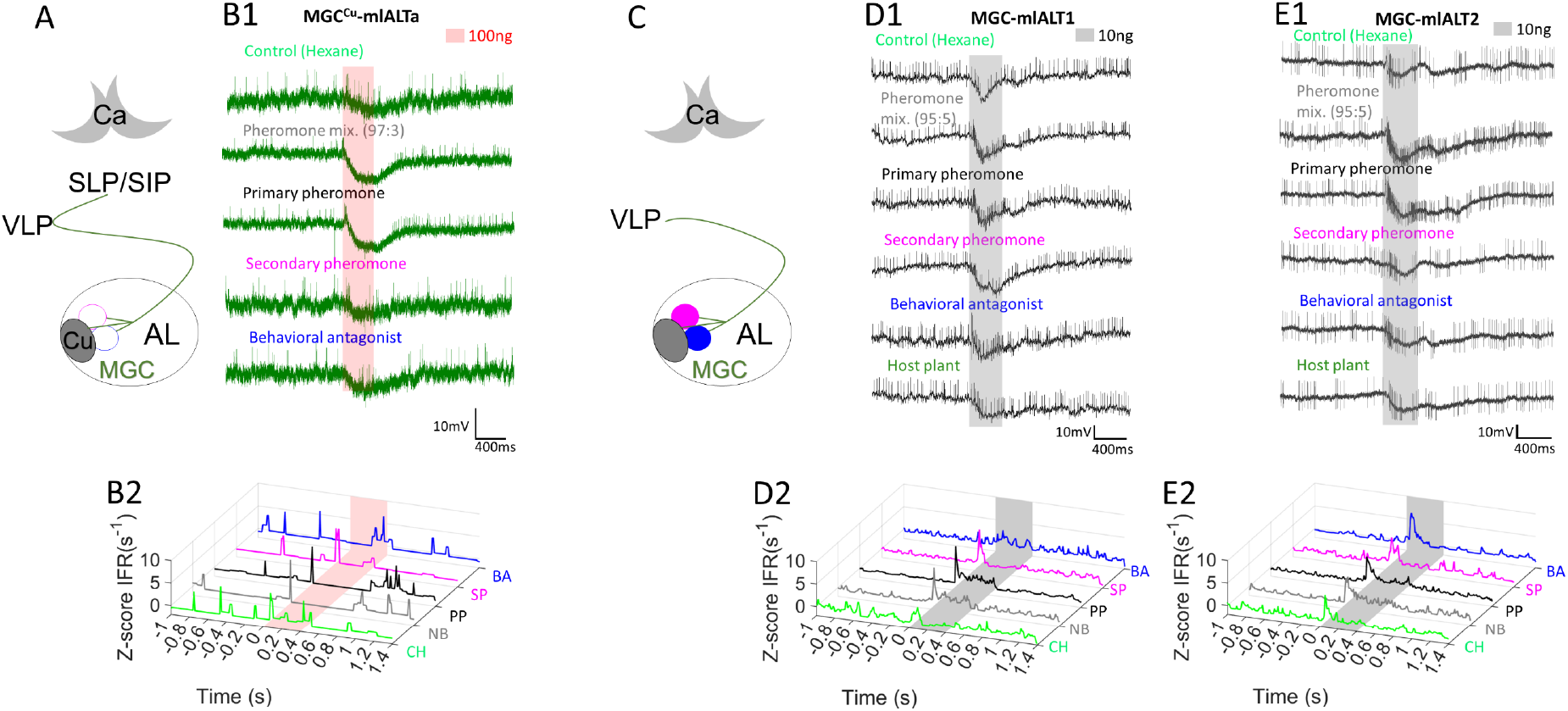
Response profiles of mlALT PNs innervating the MGC. (**A**) Schematic display of a MGC mlALT PN innervating mainly the cumulus. (**B**) PN MGC^Cu^-mlALT arborized densely in the cumulus, and had sparse dendrites in the dma, dmp, and one posterior complex glomerulus. This neuron had minor early-onset excitations followed by inhibition during stimulation with the primary pheromone (PP) and the pheromone mixture (PM). (**C)** Schematic display of a mlALT PN innervating the MGC. (**D-E**) Two multiglomerular mlALT PNs innervated all MGC units evenly, both PNs were widely tuned and displayed early-onset excitatory responses to all female-produced stimuli. Most notably, the pheromone mixture (PM) and primary pheromone (PP) elicited responses lasting throughout the entire 400 ms stimulation. In addition to the pheromone responses, MGC-mlALT2 was phasically excited by the control stimulus hexane (CH).

**Figure 5 – figure supplement 1.**
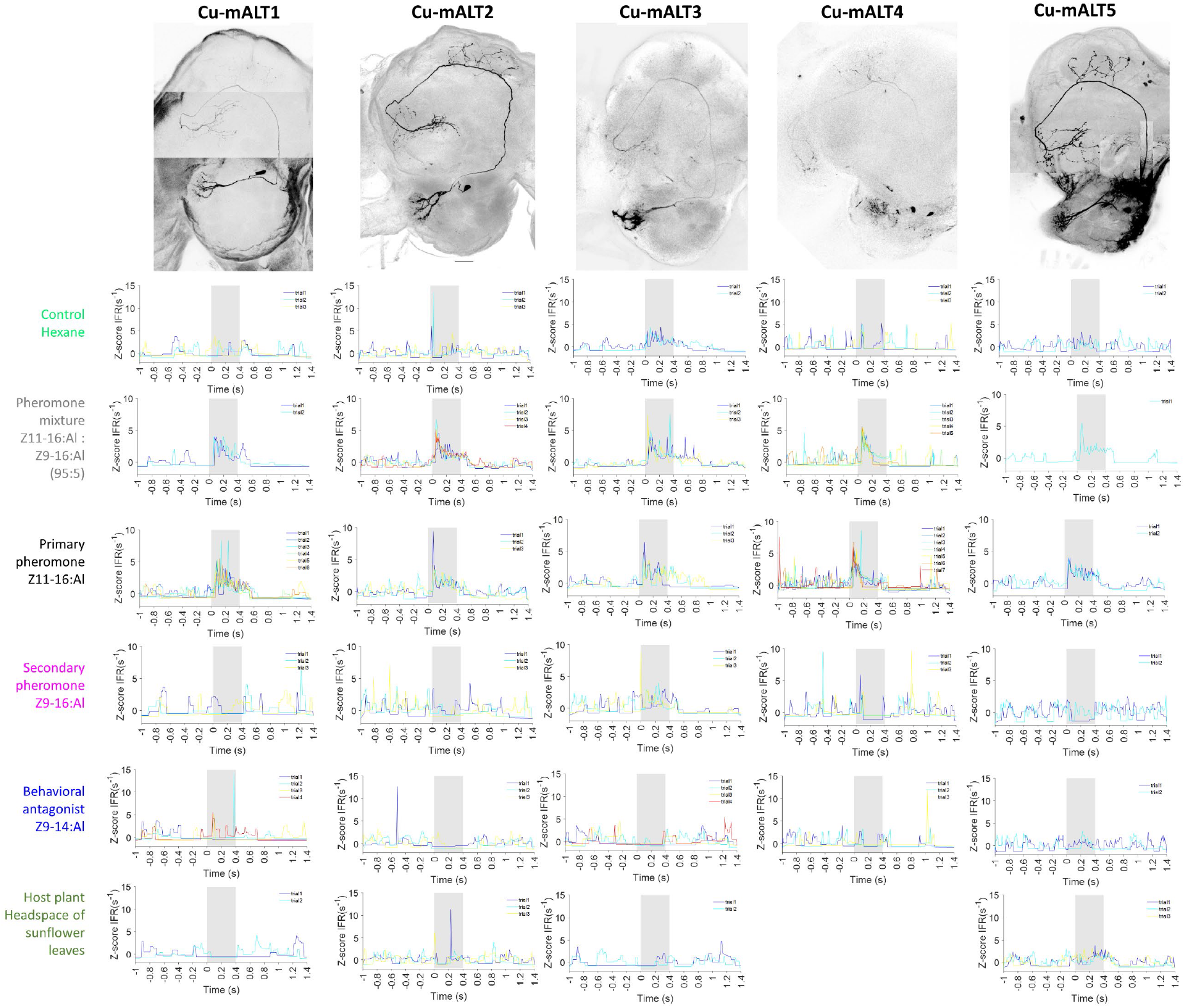
Z-scored instantaneous firing rate for each trial in cumulus PNs (Cu-mALT1-5).

**Figure 5 – figure supplement 2.**
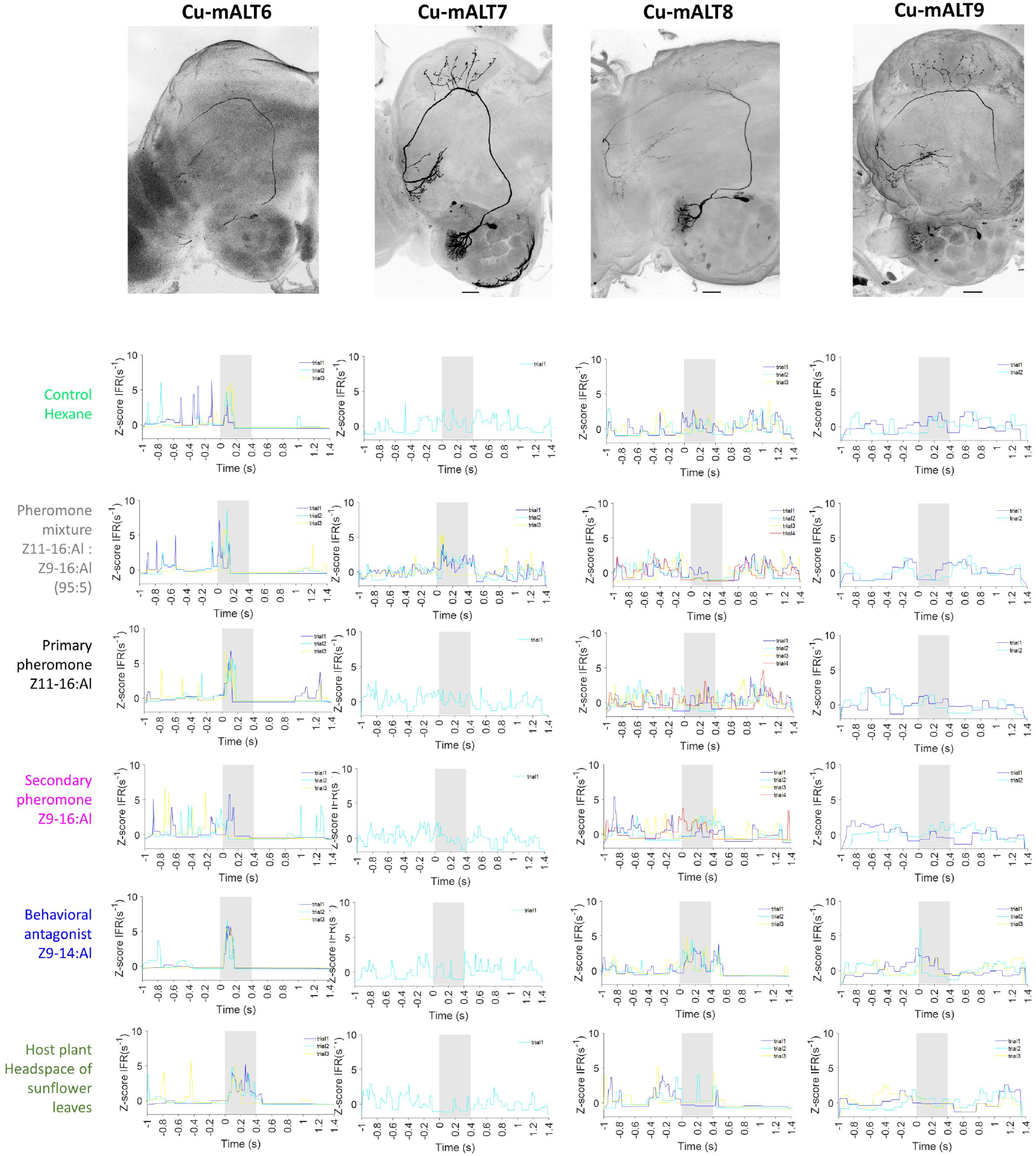
Z-scored instantaneous firing rate for each trial in cumulus PNs (Cu-mALT6-9).

**Figure 5 – figure supplement 3.**
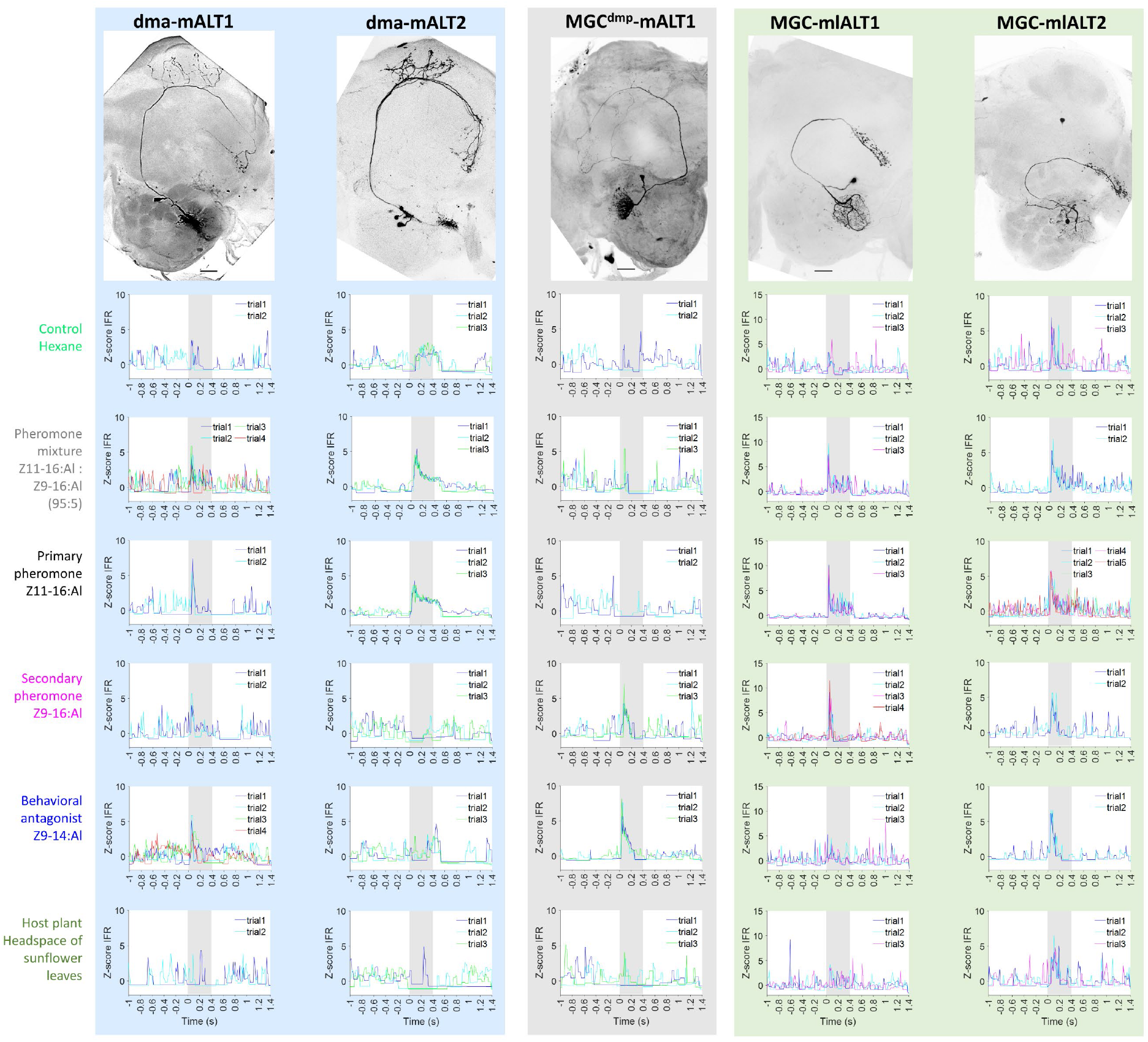
Z-scored instantaneous firing rate for each trial in dma PNs (dma-mALT1-2) and three multiglomerular MGC PNs.

**SUPPLEMENTARY TABLE S1.**
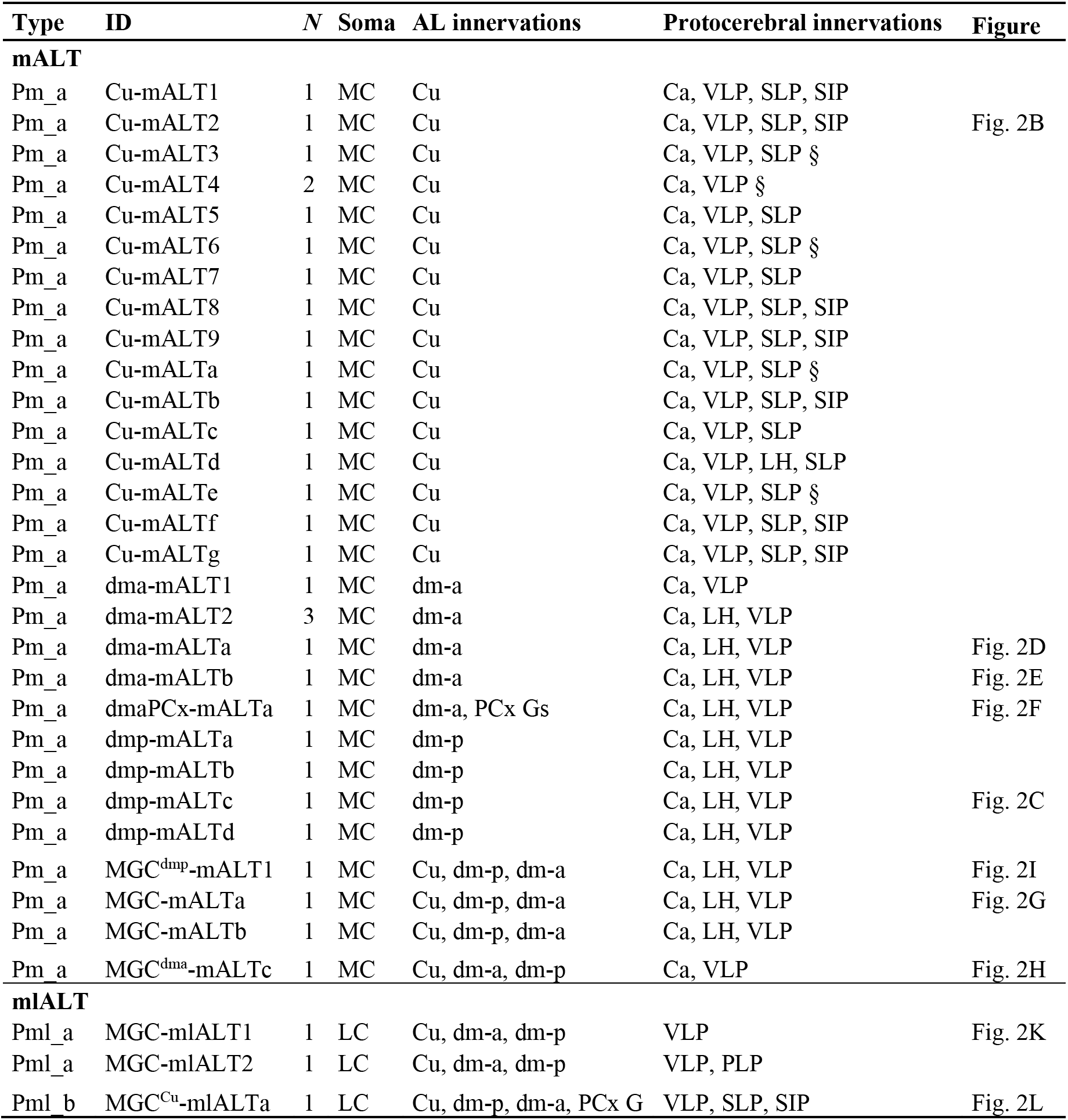
Overview of individual projection neuron morphology

